# Developmental and regulatory logic of cannabis glandular trichomes resolved by single cell multiomics

**DOI:** 10.64898/2026.05.17.725801

**Authors:** Lee J. Conneely, Matthew T. Welling, John Humphries, Lim Chee Liew, Antony Bacic, Mathew G. Lewsey

## Abstract

Glandular trichomes of *Cannabis sativa* produce specialised metabolites including cannabinoids and terpenes, yet the cellular basis of trichome development and metabolic specialisation remains unresolved. Here we reconstruct the glandular trichome lineage within the *C. sativa* inflorescence using single-cell transcriptomics and chromatin accessibility profiling. We identify a continuous developmental trajectory from the protoderm, linking bulbous, sessile, and capitate-stalked trichomes, indicating that these morphologies represent sequential stages rather than distinct trichome types. We further identify a previously undescribed neck cell that shares a developmental origin with secretory disc cells before lineage bifurcation. Integration of chromatin accessibility with transcription factor (TF) binding reveals cell state specific regulatory programs coordinating neck cell development and specialised metabolism, redox homeostasis, and secretion in mature secretory disc cells. Methyl jasmonate elicitation induces extensive transcriptional and chromatin landscape reprogramming in mature secretory disc cells and transient repression of specialised metabolic pathways. Together, these findings provide a cell-resolved regulatory framework for glandular trichome development and function, revising prevailing models of trichome identity, maturation and defence associated metabolic regulation.

## Main

Glandular trichomes are specialised epidermal micro-organs that synthesise, secrete, and store diverse bioactive metabolites, including terpenes and cannabinoids, which contribute to plant defence and have considerable pharmacological and commercial value. Increasingly, the chemistry and cell biology of glandular trichomes are attracting attention in plant synthetic biology, as these structures provide a natural platform for the heterologous production and storage of high value metabolites^1,2^. However, the cellular organisation and regulatory programs that establish and maintain this specialised secretory state remain poorly understood.

In particular, the mature *C. sativa* glandular trichome is defined by an unusually rich secretory chemistry^3^. Female inflorescences accumulate a resin-rich secretion dominated by acidic phytocannabinoids and accompanied by a chemically diverse terpene fraction comprising both monoterpenes and sesquiterpenes, together conferring much of the biological, sensory and commercial value of cannabis flowers^4,5^. These metabolites are produced by the secretory disc cells and deposited into the subcuticular storage cavity above them, tightly coupling trichome differentiation with specialised metabolic output^6^. Understanding glandular trichome development therefore requires not only a structural framework, but also an account of how this multicellular epidermal outgrowth acquires the capacity to synthesise, secrete and sustain diverse metabolites.

Transcription factors (TFs) are well established as central regulators of glandular trichome biology in better characterised systems, where factors such as SlMYC1 in tomato and multiple MYB, HD-ZIP, AP2/ERF, WRKY, and bHLH regulators in *Artemesia annua* link gland development with specialised metabolite production^7–13^. In *C. sativa,* by contrast, direct evidence for transcriptional control of glandular trichome metabolism is only beginning to accumulate. Initial support came from a promoter-focused study in which a 548-bp *THCAS* promoter fragment was used in yeast one-hybrid and transient reporter assays, identifying CsAP2L1 as a positive regulator of *THCAS* promoter activity and CsWRKY1 and CsMYB1 as putative negative regulators^14^. Subsequent work linked additional cannabis TFs to glandular trichome development, including CsMIXTA, which increased glandular trichome density in heterologous tobacco assays, and the jasmonate-responsive bHLH factor CsMYC4, which was implicated in methyl-jasmonate (MeJA) mediated glandular trichome formation^15,16^. Most recently, CsSRS116 has been identified as a glandular trichome associated factor whose target genes were enriched for functions related to growth and development^17^. Together these studies demonstrate that we are beginning to understand the transcriptional machinery coordinating cannabis glandular trichome differentiation and secretory function, but that much remains to be discovered compared with tomato or *A. annua*.

Recent studies have begun to refine the developmental framework of *C. sativa* glandular trichomes. A revised model proposes that sessile and capitate stalked trichomes represent sequential stages along a single developmental trajectory rather than distinct morphological classes, with sessile trichomes functioning as pre-stalked intermediates^18^. Additional work has revealed that mature secretory disc cells operate as a coordinated multicellular unit connected by cytoplasmic bridges through adjacent cell wall perforations, consistent with a model of a polarised secretory “supercell”, and that extensive cell wall remodelling drives delamination and formation of the secretory cavity during gland maturation^19,20^. Complementary proteomic, transcriptomic, and epigenomic datasets generated through physical enrichment of trichomes have further advanced our understanding of the specialised metabolism underpinning these micro-organs^21–23^. Recent studies have also explored the transcriptional responses of *C. sativa* inflorescences to methyl jasmonate defence elicitation, highlighting the roles of defence signalling in regulating trichome development and specialised metabolism^16,24,25^. However, the cellular organisation, developmental transitions, and regulatory programs governing glandular trichome development and defence responses remain incompletely resolved.

To resolve the cellular and regulatory basis of glandular trichome development, we investigated the histological, transcriptional, and chromatin landscapes of the *C. sativa* inflorescence. We combined synchrotron micro-computed tomography (micro-CT), single-nucleus multiomic profiling and DNA affinity purification sequencing with classical RNA *in situ* hybridization to reconstruct the glandular trichome developmental lineage and its associated regulatory programs. We identify a single developmental trajectory for glandular trichomes that supports previous models proposing sessile trichomes as pre-stalked intermediates of capitate stalked trichome development, and from which we further propose that bulbous trichomes represent an earlier intermediate stage preceding this transition^18^. We also report the discovery and validation of previously undescribed glandular trichome neck cells, a transcriptionally distinct cell type that shares a developmental origin with secretory disc cells. We uncover parallel state-specific TF modules operating within neck and disc cell lineages, including a novel transcriptional regulator of cannabinoid and specialised metabolism biosynthetic pathways and secretory processes in mature disc cells. Lastly, we examine the transcriptional and chromatin landscape rewiring in mature secretory disc cells following treatment with the defence hormone MeJA. Together, these analyses revise the prevailing model of cannabis glandular trichome development by resolving bulbous, pre-stalked, and capitate stalked trichome not as independent trichome types, but as discrete developmental states within a single epidermal differentiation trajectory.

## Results

### Structural organization of the *Cannabis sativa* inflorescence and trichome development

We first defined the structure of the *C. sativa* inflorescence (Fig. 1a) and the developmental context of its glandular trichomes, to support downstream single-nucleus multiomics and histological analyses. A longitudinal section imaged using micro-computed tomography (micro-CT) revealed a compound raceme architecture with densely imbricated bracts enclosing individual flowers (Fig. 1b). Transverse sections confirmed the bicarpellate gynoecium, reduced leaf structures and prominent stigmatic tissues (Fig. 1c). The developing flower lacked trichomes on the gynoecium epidermis and was instead enclosed by three imbricate bracts enriched for both glandular and non-glandular trichomes. Trichome composition varied by bract layer: the innermost bract, adjacent to the gynoecium, was enriched in cystolithic hairs (non-glandular trichomes), whereas the two outer bracts were enriched in glandular trichomes (Fig. 1d).

**Fig. 1.**
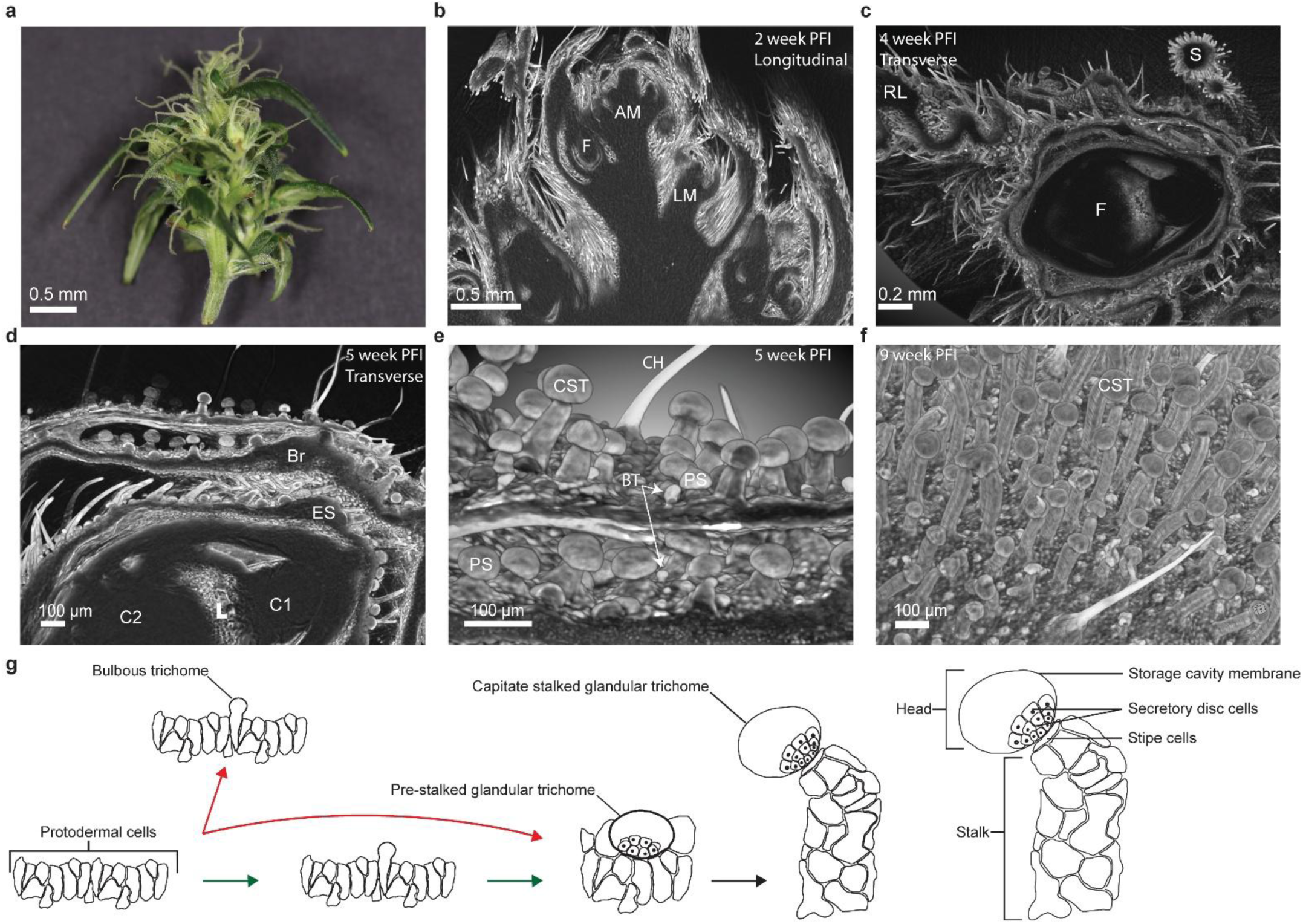
| Structural organization of the *C. sativa* inflorescence and trichome development. **a**, Longitudinal view of a *C. sativa* inflorescence 4 weeks post floral induction (PFI). **b**, Longitudinal micro-CT section of a 2-week PFI inflorescence showing the apical meristem (AM), lateral meristem (LM) and developing flower (F). **c**, Transverse micro-CT section of a 4-week PFI inflorescence showing flower (F), reduced leaf (RL) and stigma (S). **d**, Transverse micro-CT section of a single flower from a 5-week PFI inflorescence showing three trichome-rich imbricate layers surrounding the gynoecium; outer bract layers (BR) are enriched in glandular trichomes, whereas the inner early sepal (ES) is enriched in cystolithic hairs. The locule (L) and both small (C1) and large (C2) carpels of the gynoecium are visible. **e**, Micro-CT image of the bract epidermal surface from a 5-week PFI inflorescence showing a heterogenous trichome population comprising bulbous trichomes (BT), pre-stalked trichomes (PS), capitate-stalked glandular trichomes (CST) and cystolithic hairs (CH). **f**, Micro-CT image of the bract epidermal surface from a 9-week PFI inflorescence showing CSTs as the predominant glandular trichome morphotype, with elongated stalks. **g**, Schematic of two alternative models of *C. sativa* glandular trichome development from the protoderm. In model 1, bulbous and capitate-stalked glandular trichomes arise from parallel developmental trajectories; in model 2, bulbous trichome represent an early developmental morphotype preceding capitate-stalked glandular trichome development.

We determined that four major trichome morphotypes were present using micro-CT imaging at two, four, five, and nine weeks post-floral induction (PFI): cystolithic hairs, bulbous trichomes, pre-stalked trichomes (historically termed sessile) and capitate stalked glandular trichomes (Fig. 1e). Notably, the transition from pre-stalked to capitate stalked forms was not synchronous across the population. Instead, we observed a continuum of intermediate morphologies, indicating that trichome maturation is not coupled to overall inflorescence development during early and mid-stages. By nine-week PFI, trichome populations were largely uniform, with capitate stalked glandular trichome predominating (Fig. 1f).

Based on these observations and prevailing models of glandular trichome ontogeny, we propose a developmental framework in which glandular trichome initiation begins in protodermal cells, and pre-stalked (historically termed sessile) trichomes are an intermediate stage preceding capitate stalked glandular trichomes^18,26,27^ (Fig. 1g). However, whether a discrete transitional state links protodermal cells to morphologically distinct pre-stalked glandular trichomes remains unresolved. Historically classified as a separate trichome type, we supposed that bulbous trichomes may be a microscopy-defined class that may instead represent a transitional developmental state, rather than a separate trichome class.

### Construction and cell-type annotation of a *Cannabis sativa* inflorescence single-nucleus multiomic atlas

We next sought to define cell type-resolved transcriptomic and chromatin accessibility landscapes of *C. sativa* glandular trichomes, to enable reconstruction of their developmental trajectory from protodermal origins. We applied the 10x Genomics Multiome approach, which generates single-nucleus RNA-seq (snRNA-seq) and assay for transposase accessible chromatin (snATAC-seq) data simultaneously from individual nuclei, to isolated nuclei from 6-week PFI inflorescence under control conditions and two-hours after foliar spray treatment with 1 mM MeJA. This was enabled by developing a bench-top nuclei isolation workflow compatible with 10x Genomics Multiome chemistry and avoiding DNA-binding dyes, many of which are contraindicated for snATAC and snMultiome assays due to detrimental effects on data quality and UMI recovery (Fig. 2a)^28^.

**Fig. 2.**
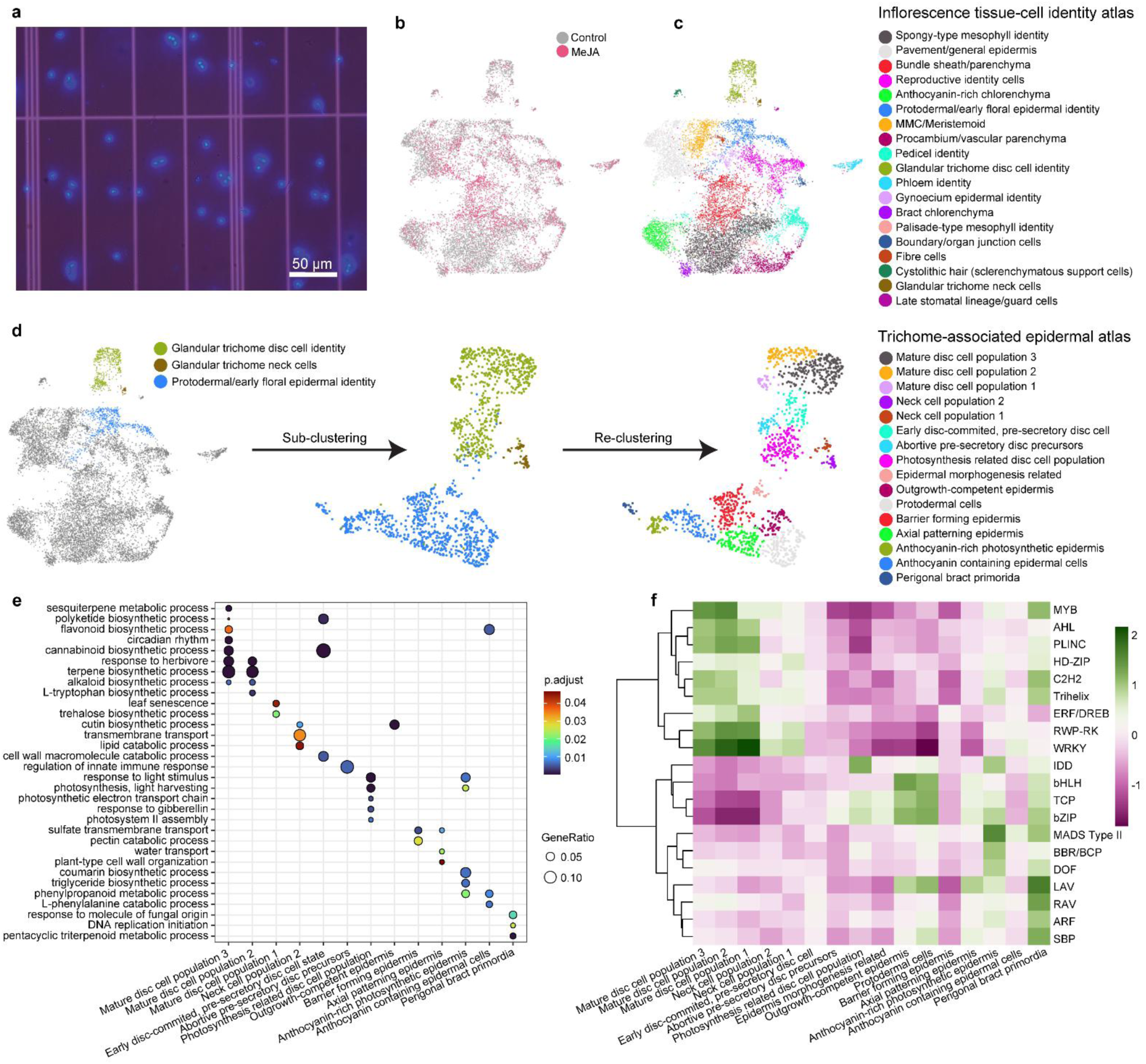
| Construction and annotation of a *C. sativa* trichome-associated epidermal single-nucleus multiomic atlas. **a**, A DAPI-stained nuclei isolated from a 6-week PFI *C. sativa* inflorescence. **b**, Weighted nearest neighbour (WNN) UMAP of inflorescence nuclei coloured by control (grey) and MeJA-treated (pink) conditions. **c**, WNN UMAP showing cluster resolution and cell-type annotation of inflorescence nuclei. **d**, Subclustering of trichome-associated epidermal cells, showing refined cell types and transcriptional states. **e**, Gene ontology (biological process) enrichment across clusters based on the top-ranked marker genes (logFC; adjusted *P* < 0.05) using PANNZER2-derived annotations. **f**, Heatmap of chromVAR-derived transcription factor motif family enrichment (JASPAR 2024) across cell types, shown as Z-score-scaled accessibility.

Doublet and plastid-aware filtering was applied to the data yielding strong transcription start site (TSS) enrichment of ATAC counts, evidence of preserved chromatin structure, as well as comparable quality control (QC) metrics across samples for both ATAC and RNA components (Extended Data Fig. 1a-d). Following quality control, 11,926 high-quality nuclei were retained across two biological replicates for both control and MeJA treatment conditions (Fig. 2b).

We next defined transcriptionally distinct cell types present in the *C. sativa* inflorescence by clustering of nuclei. Harmony integration between samples was applied prior to multimodal (snRNA+snATAC-seq) analysis to mitigate residual sample-associated variation in RNA embeddings^29^. This increased neighbourhood mixing (iLISI) from 1.72 to 2.46 and indicated improved cross-sample alignment. Major cell identities were resolved, whilst avoiding over-fragmentation of transcriptionally distinct cell states, by clustering nuclei using a multimodal weighted nearest-neighbour (WNN) graph that integrated transcriptomic and chromatin accessibility modalities^30^. Nineteen discrete clusters were resolved, which we annotated with the key aim of identifying clusters that represent glandular trichome associated cells (Fig. 2c). Cell-type identities were assigned using literature-curated marker genes and gene ontology (GO) enrichment analysis as ground truth single-cell datasets do not exist for *C. sativa* inflorescence (Extended Data Fig. 1e, and Supplementary Table 1a-f).

Three adjacent clusters were inferred to comprise trichome-associated epidermal cells: protodermal/early floral epidermis, glandular trichome neck cells, and glandular trichome disc cells. Annotation of the protodermal/early floral epidermis population was based upon the enrichment of *PDF1* (LOC115699665)^31^. Glandular trichome neck cells were identified using *MYB36* (LOC115704480), an ortholog of the Casparian strip master regulator recently identified in *Cucumis sativa* trichome neck cells, as well as a cluster specific Casparian strip gene *CASPL1C1* (LOC115709649) (Fig. 2c, Supplementary Table 1a,f)^32^. Glandular trichome disc cells were collectively defined by expression of known markers including delta-12 oleate desaturase (*FADX;* LOC115719251), geranyl diphosphate synthase small subunit (*GPPSss;* LOC115725388) and cannabidiolic acid synthase-like (*CBDAS*-like; LOC115696884)^18,22,33^.

### Individual glandular trichome cell types exhibit multiple transcriptional cell states and chromatin landscapes

We performed a focused analysis to resolve the trichome-associated epidermal cells in greater detail. We isolated just the protodermal/early floral epidermal, glandular trichome neck cells, and glandular trichome disc cell identity clusters, then re-integrated and re-clustered them. This increased neighbourhood mixing (iLISI 2.28 to 3.06), indicating improved cross-sample alignment. Sixteen subclusters were resolved by this approach, which represented transcriptionally distinct cell states (Fig. 2d). Cell-type and state identities were assigned to these clusters using the following criteria: GO term enrichment analysis (BP, CC and MF) with PANNZER2-annotated cs10 (GCA_900626175.2) proteins, accessible TF-binding motif enrichment, examination of literature-curated marker genes, and experimental validation by RNA *in situ* hybridization of marker transcripts (Fig. 2e, f, Fig. 3, Extended Data Fig. 1f, Extended Data Fig. 2-4, and Supplementary Table 2a-j,3)^34–37^.

**Fig. 3.**
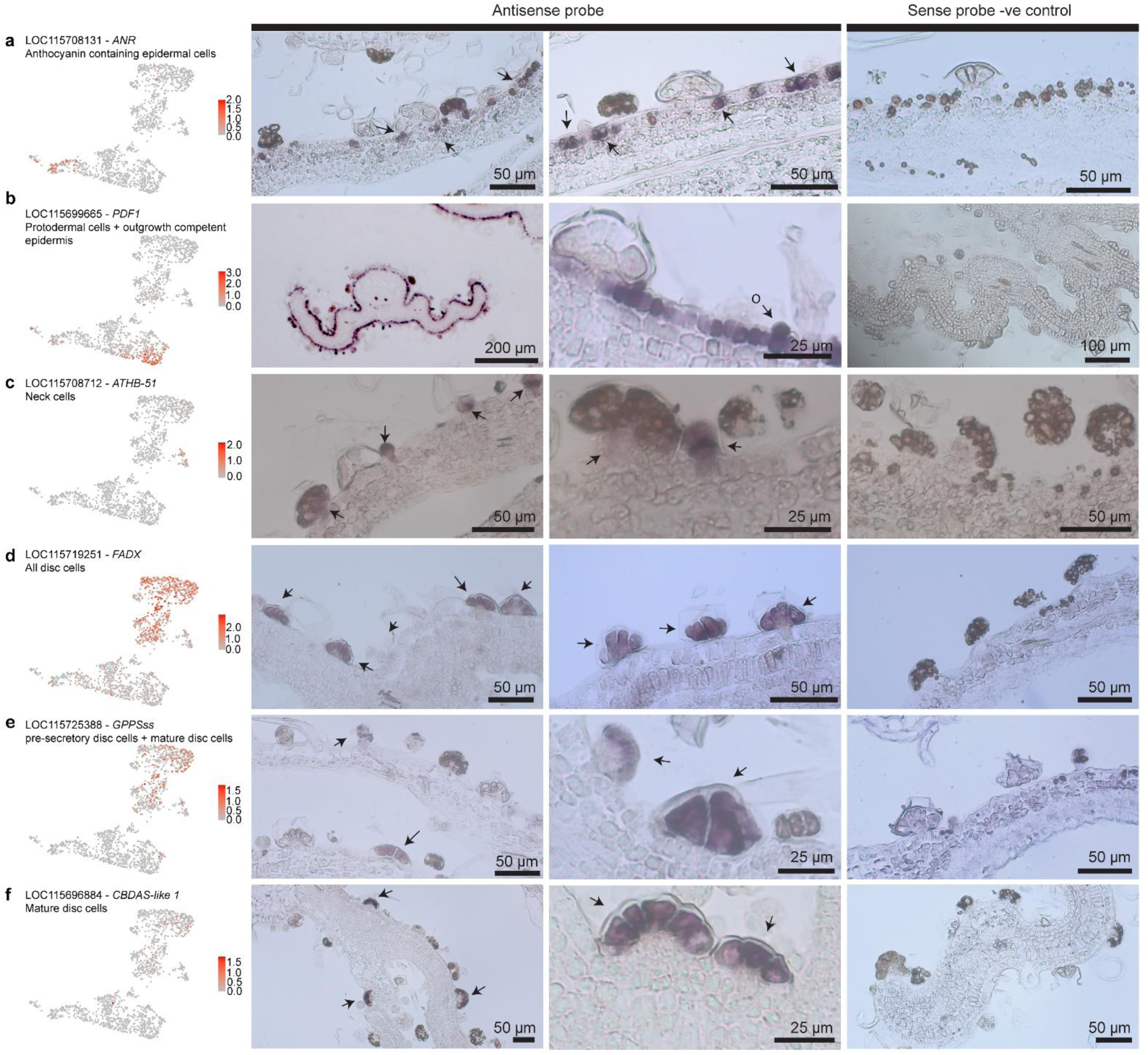
| Validation of cell-type assignments by RNA *in situ* hybridization. RNA *in situ* hybridization on sections of 5-week post floral induction (PFI) *C. sativa* inflorescences was used to validate cell-type assignments in the trichome-associated epidermal atlas. Antisense probe signal (purple) was interpreted relative to the corresponding sense-probe negative controls. **a**, *ANR* (LOC115708131) transcripts in anthocyanin-containing epidermal cells. Signal marks pigmented epidermal cells and distinguishes this population from adjacent protodermal cells. **b**, *PDF1* (LOC115699665) transcripts in the outer epidermal layer. Signal is discontinuous and patchy, with enrichment in protruding, outgrowth competent epidermal cells (O). **c**, *ATHB-51* (LOC115708712) transcripts in glandular trichome neck cells. Signal appears as a distinct band at the neck region, subtending early pre-secretory disc cells, and remains detectable during trichome development, although with reduced intensity at later stages. **d**, *FADX* (LOC115719251) transcripts mark glandular trichome disc cells. Residual trichome resin was occasionally retained after clearing but did not preclude assignment of probe signal. **e**, *GPPSss* (LOC115725388) transcripts in early disc-committed, pre-secretory disc and mature disc cells. **f**, *CBDAS-like* (LOC115696884) transcripts restricted to mature glandular trichome disc cells.

Cells with protodermal/early floral epidermal identity resolved into eight subclusters, indicative of distinct cell states (Fig. 2d). Five of these subclusters were from adjacent developmental programs (i.e. not trichomes) related to early bract development and pigment containing epidermal cell development. These included a ‘perigonal bract primordia’ population that was enriched for floral homeotic genes *AG* and *AGL19* and MADS-box class II motif accessibility and motif-target gene modules enriched for plant organ formation, an ‘axial patterning epidermis’ cluster marked by the sulfate transporter *SULTR3-5* (LOC115703621) and the axial regulator *YABBY5* (LOC115724567) in developing bracts, a ‘barrier forming epidermis’ cluster enriched for *CASP* genes and *CalS10* (Supplementary Table 2a-i and Extended data Fig. 2-4)^38–41^. Two closely-related cell types, ‘anthocyanin-rich photosynthetic epidermis’ and ‘anthocyanin containing epidermal cells’, were characterised by enrichment of flavonoid and phenylpropanoid GO terms, and restricted expression of the anthocyanin biosynthetic enzyme anthocyanidin reductase *ANTR* (LOC115708131) in visibly pigmented, flat, epidermal cells, consistent with pigment containing, epidermal cells and distinguishable by the enrichment of photosynthesis related biology in ‘anthocyanin-rich photosynthetic epidermis’ (Fig. 3a and Supplementary Table 2a-i)^42^.

The remaining three subclusters with protodermal cell/floral cell identity were within the early protoderm-trichome lineage (Fig. 2d, 3b, Extended Data Fig. 2, and Supplementary Table 2). There were canonical ‘protodermal cells’, which expressed *PDF1* (LOC115699665) and further characterised by the enrichment of class II Teosinte Branched1/Cycloidea/Proliferating cell factor (TCP) regulators including *TCP4, TCP2,* and accessible chromatin enriched for TCP binding motifs, which are consistent with a regulatory program that coordinates the proliferation to differentiation ratio and restricts excessive trichome initiation^43,44^. Continuous with the protodermal cells was a putative ‘outgrowth competent epidermis’ cell state, enriched for SCF- type E3 factor (LOC115698583), an ortholog of *At2g36090* linked to endoreduplication and outgrowth competency in early trichome development in *A. thaliana*. Interestingly, MYC4 (MA0569.2) motif accessibility was enriched in this cell type although MYC4 transcripts were not detected. CsMYC4 was recently identified as a key early regulator in a MeJA mediated trichome development pathway in *C. sativa*. Taken together these appear likely to describe a cell-state chromatin landscape that is primed for trichome development with capacity to integrate defensive hormonal cues^16,45^. There was also an ‘epidermal morphogenesis’ cluster enriched for genes linked to trichome density, cell morphogenesis, and cell polarisation, including *NAC25* (LOC115697858), *SRF3* (LOC115697792), and *ROPGEF12* (LOC115704879) (Supplementary Table 2a-i and Extended Data Fig. 2)^46–48^.

The neck cells partitioned into two states, which we provisionally termed neck cell populations 1 and 2. Both neck cell populations expressed the *ATHB-51* TF (LOC115708712), which exhibited spatially restricted expression in the neck cells subtending secretory disc cells (Fig. 3c). Neck cell population 1 was enriched for borate-transport related processes, the early neck program regulator *MYB36* (LOC115704480), a transcriptional regulator of suberin biosynthesis *MYB53* (LOC115715904), and accessible HD-ZIP class I binding motifs including ATHB-51 (MA0952.1), the ATHB-51 motif was previously shown to be enriched in putative enhancer elements specific to *C. sativa* glandular trichome isolates when compared with leaf and stalk bulk tissues datasets^32,49–51^. This suggested epidermal cell fate determination and trichome development programs were active in this neck population. Neck cell population 2 was enriched for secondary cell wall metabolism, Casparian strip genes, and the suberin biosynthesis marker *HHT1* (LOC115699431)^52^.

Subclustering also resolved a continuum of transcriptionally distinct disc cell states, spanning a ‘photosynthesis-related disc cell’, ‘early disc-committed pre-secretory cell’, ‘abortive pre-secretory cell’, and three mature secretory cell populations (Fig. 2d). Collectively, disc cell identity was confirmed by RNA *in situ* hybridization of the marker transcript *FADX* (LOC115719251), whilst the markers *GPPSss* (LOC115725388) and *CBDAS-like 1* (LOC115696884) characterised the early and mature populations (Fig. 3d-f)^18,21,22^. We note that residual resin was visible in disc cells after clearing of sections, but antisense probe signal remained clearly detectable in resin-containing disc cells and no comparable signal was observed in sense-probe controls, indicating that probe sensitivity and specificity were not compromised.

The photosynthesis-related disc cell state was enriched for plastid- and peroxisome-associated CC ontologies, expressed MEP pathway gene DXR (LOC115709371) and the early cannabinoid pathway gene *OAC* (LOC115723437), and was enriched for the accessibility of JKD/MGP and IDD-family motifs together with high expression of *IDD4, IDD5,* and *IDD7* homologues, consistent with active programs supporting photosynthesis, starch metabolism, and tissue differentiation (Supplementary Table 2a-i)^53–55^. Abortive pre-secretory cells similarly expressed genes upstream of cannabinoid and terpenoid biosynthesis, but were distinguished by features associated with cellular stress and plant immunity, including *PUB3* (LOC115718652), the autophagy marker *Beclin-1* (LOC115714331), and defence response to symbiont and innate immune response BPs^56,57^. Early disc-committed pre-secretory disc cells were highly enriched for expression of genes associated with early cannabinoid biosynthesis, polyketide biosynthesis, and several cell division related BPs. These observations suggest disc-cell fate commitment and active cell division to generate the canonical ∼12-16 disc cells found in *C. sativa* mature glandular trichomes.

Mature disc cell states (mature disc cell populations 1, 2, and 3) exhibited enriched expression of terminal cannabinoid and terpenoid biosynthetic genes (Fig. 2, Extended data Fig. 2, 3 and Supplementary Table 2a-c). WRKY motif accessibility followed a gradient of accessibility, from highest in mature disc cell population 1 to lowest in population 3. A similar gradient was observed in expression of WRKY-motif target genes, which were enriched for abiotic and biotic stress response, salicylic acid, and phenylpropanoid metabolism related terms consistent with a defence associated regulatory program^58^. Most interestingly, in the snRNA-seq data we observed enrichment of the trehalose biosynthetic pathway term in marker gene GO analysis, including two putative TPS genes and a TPP, which suggested mature disc cell population 1 engages in active sugar signalling circuitry associated with carbon sufficiency, sink establishment and metabolic priming^59,60^. The trichome development factor *HDG11* (LOC115714770) was highly enriched in mature disc cell population 1 and to a lesser extent in populations 2 and 3. The ortholog of this gene in *A. thaliana* is a trichome development stage marker, indicating mature disc cell population 1 may be a relatively earlier developmental stage of disc cell development compared with populations 2 and 3^61^.

MYB motif accessibility was highly enriched in mature disc cell populations 2 and 3, and MYB-motif target-gene expression mirrored this pattern and comprised of isoprenoid-, terpene-, terpenoid-, and sesquiterpene-related ontology terms (Supplementary Table 2c-g). Notably, the *WRI1* (LOC115706982) master regulator of plant oil biosynthesis was a marker for both mature disc cell populations 2 and 3, indicating fatty acid biosynthesis is likely enhanced in these cell types^62^. Mature disc cell population 2 was highly enriched in diverse transporters including A,B,C,D,F and G class ABC transporters, including well-known specialised metabolite transporters *PDR1* (LOC115705239; LOC115705243) and flavonoid transporter ABCC2 (LOC115697398)^63,64^. Mature disc cell population 3 was highly enriched for myrcene and α-humulene transcripts, moreover a bulk time-course analysis of *C. sativa* trichome head metabolites revealed myrcene and α-humulene concentration was highest during the final plant development time point T8, which suggested that mature disc cell population 3 may be the more mature disc cell state of the three mature disc cell populations^65^. Taken together these data indicated that specialised metabolism, fatty acid biosynthesis, and secretion are active in mature disc cell populations 2 and 3 and are distinguished by discreet changes in specialised metabolism (Supplementary Table 2a-i).

The position of the photosynthesis-related disc cell population within the UMAP projection suggested that these cells may be an intermediate transcriptional state between protodermal epidermis and differentiated secretory disc cells. Notably, photosynthesis-associated GO terms were elevated in the photosynthesis related disc cell population and were reduced in pre-secretory and mature disc cells, although plastid membrane and chloroplast envelope CC annotations were retained (Supplementary Table 2c,d). This aligns with the known absence of photosynthetically active plastids in *C. sativa* glandular trichomes and their differentiation into specialised non-photosynthetic leucoplasts that support MEP-derived isoprenoid biosynthesis ^19,21,66^. We therefore propose that initial disc cell maturation may involve transient activation of chloroplast biogenesis and photosynthesis, followed by plastid specialisation toward high-capacity isoprenoid production that fuels terpene and cannabinoid biosynthesis.

### Glandular trichomes originate from a single developmental trajectory with a neck-disc cell bifurcation

We next reconstructed the developmental trajectory of glandular trichomes so that we might better understand the structure of this process and how the three trichome morphotypes (bulbous, pre-stalked, and capitate-stalked) relate to one-another. To do this we reconstructed a pseudotime trajectory to represent their development, rooted in the protodermal cells as known progenitors and incorporating the 11 glandular trichome-annotated clusters (Fig. 2d, 4, Extended Data Fig. 5). All other clusters were excluded. The reconstructed trajectory proceeded through transitional epidermal states to terminal secretory disc cells, with a single bifurcation point between neck and disc cell fate (Fig. 4a, b).

**Fig. 4.**
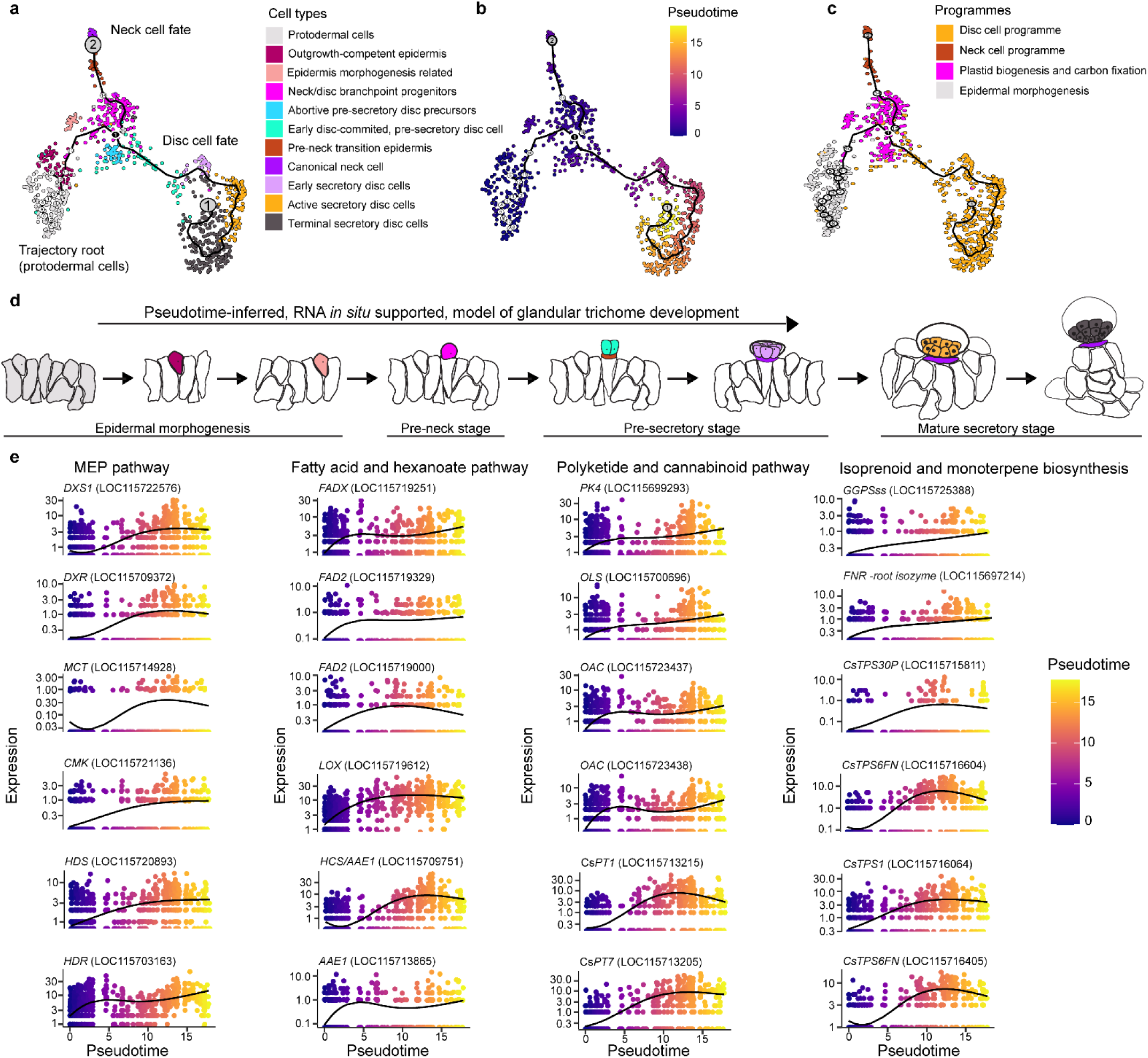
| A single developmental trajectory links protodermal, disc and neck cell states in glandular trichomes. **a**, Monocle 3 trajectory analysis of trichome-lineage cells rooted in protodermal cells, revealing a continuous developmental path towards terminal secretory disc cells with a bifurcation towards neck cell fate at the photosynthesis-related disc-cell population (neck/disc branchpoint progenitors). **b**, Pseudotime projection of trichome-lineage cells. Mature disc cells span a broad interval of pseudotime, consistent with substantial heterogeneity within the terminal branch, whereas neck cells form a narrow branch with limited pseudotime dispersion. **c**, Gene co-expression modules identified from genes with trajectory-structured expression (Moran’s *I* > 0.1, adjusted *q* < 0.01). Cells are coloured by trajectory-informed module co-expression scores. **d**, Schematic model of glandular trichome development inferred from pseudotime analysis and supported by RNA in situ hybridization, showing a single developmental trajectory encompassing all trichome morphotypes. **e**, Literature-curated genes associated with trichome specialised metabolism and showing trajectory-structured (Moran’s *I* > 0.1, adjusted *q* < 0.01), plotted across pseudotime. Genes from the MEP, fatty acid and hexanoate, terminal cannabinoid, isoprenoid and monoterpenoid pathways increase strongly from pseudotime 5, corresponding to mature disc cells and early cannabinoid precursor biosynthesis including fatty acid biosynthesis, polyketide, and early cannabinoid biosynthetic genes are induced in early pseudotime points corresponding with the early disc-committed, pre-secretory disc cells.

The reconstructed trajectory provided notable insight into glandular trichome development. Foremost, no bulbous trichome-specific branch, nor a terminal bulbous trichome state, was identified within the trajectory (Fig. 4a). This argues against the presence of a developmentally discrete bulbous glandular trichome class and instead supports a developmental model in which bulbous trichomes are an intermediate morphogenetic stage within the capitate-stalked glandular trichome ontogeny. Consistent with this model, the neck cell marker transcript *ATHB-51* (LOC115708712) was expressed strongly and in a spatially restricted domain within early stage glandular trichomes that had bulbous morphology, as observed by RNA *in situ* hybridization (Fig. 3c). The *ATHB-51* signal was most pronounced during early outgrowth and progressively diminished in more mature glandular trichomes exhibiting pigmented secretory disc cells and expanded extracellular storage cavities. This observation correlated with the expression pattern along the pseudotime trajectory (Extended Data Fig. 5).

The trajectory also indicated that a branchpoint exists that bifurcates neck and disc cell fates, suggesting an asymmetric cell fate decision that establishes structural (neck) and secretory (disc) compartments (Fig. 4a, b). The cells at this branchpoint on the trajectory had been initially annotated as a ‘photosynthesis related disc cell population’ (Fig. 2d), so we henceforth renamed them neck/disc branchpoint progenitors (Fig. 4a). The neck cell fate then proceeded through the cluster originally termed neck cell population 1, which we renamed as ‘pre-neck transition epidermis’. This population was shown earlier to be enriched in orthologous early neck cell regulators, which supports our trajectory analysis wherein this cell type is not terminally differentiated. The fate terminated in neck cell population 2, consequently renamed as canonical neck cells. This was interpreted to be a terminally differentiated neck cell state, comparatively depleted of early neck regulators and enriched for suberin, lignin, and phenylpropanoid biosynthetic pathway genes (Fig. 4a,b).

The disc cell fate proceeded through early disc-committed pre-secretory cell, mature disc cell populations 1, 2 and 3 sequentially. Of the three mature disc cell populations mature disc cell population 1 was earliest on the pseudotime trajectory and consequently we renamed it early secretory disc cells. These were defined by expression of canonical disc related genes but clearly distinguished by enrichment of accessible WRKY TF binding sites (Supplemental Table 2a,b,f). Mature disc cell population 2 was renamed active secretory disc cells, based upon the enrichment of accessible binding sites for MYB TFs known to regulate specialised metabolism and high expression of terminal biosynthetic pathway genes, including cannabinoids and monoterpenes and diverse classes of specialised metabolite transporters (Supplemental Table 2a,b,f). Mature disc cell population 3 was at the end of the reconstructed trajectory and so renamed as terminal secretory disc cells. These exhibited a shift toward sesquiterpene metabolism comparatively.

We characterised the active gene expression programs during the glandular trichome developmental trajectory by modelling gene co-expression then related them to the functions of the different glandular trichome cell types. Four modules of co-expressed genes were identified along the trajectory (Fig. 4c, Extended Data Fig. 6a-c, Supplementary Table 4a,b). One module, which we termed the disc cell program, was comprised of the early disc-committed, pre-secretory disc cell, early secretory, active secretory and terminal secretory disc cells. Genes within this module were enriched for ontology terms associated with monoterpene, sesquiterpene, and specialised metabolism (Supplementary Table 4c-e). The module that we termed the neck cell program was composed of both pre-neck transition epidermis and canonical neck cells, and was enriched for lignin metabolism and biosynthesis ontologies (Supplementary Table 4c-e). We also identified a module that we termed plastid biogenesis and carbon fixation expressed in the epidermal morphogenesis related, neck/disc branchpoint progenitors, and abortive pre-secretory disc cells. Genes within this module were associated with photosynthetic activity (Supplementary Table 4c-e). The final module we termed epidermal morphogenesis and was expressed in the protodermal and outgrowth-competent epidermal cells. This module was enriched for trichome differentiation and cell morphogenesis ontologies (Supplementary Table 4c-e). Together, trajectory inferred co-expression modules highlight the major transcriptomic changes that occur during trichome development, from multipotent protodermal cells to terminally differentiated and metabolically active mature disc cells. We then made a schematic representation of the major developmental transitions occurring during capitate stalked glandular trichome development interpreted from the combined histological and trajectory analysis in this study (Fig. 4d).

We next examined how cannabinoid biosynthesis and specialized metabolism were co-ordinated across cell-types during glandular trichome development by identifying genes with structured expression patterns (Moran’s *I* spatial autocorrelation > 0.1, q < 0.01) along the reconstructed developmental trajectory (Fig. 4e, Extended Data Fig. 5b, and Supplemental Table 4a)^67^. MEP pathway genes were progressively upregulated from early disc committed, pre-secretory disc cells and maintained through early, active and terminal secretory states, consistent with a gradual increase of isoprenoid supply. In contrast, fatty acid and hexanoate-associated genes exhibited gene-specific kinetics rather than a uniform program. For example, *FADX* (LOC115719251) and *FAD2* (LOC115719329) were initially upregulated early, with expression progressively increasing toward later disc cell states, whereas a second *FAD2* (LOC115719000) was upregulated early but then decreased in expression as the trajectory progressed^22,68^.

Downstream oxylipin/ green leaf volatile-related enzymes (*LOX*; LOC115719612, *HPL*; LOC115698766) and hexanoate activation (*HCS/AAE1*; LOC115709751) were expressed most highly in intermediate-to-late disc states, consistent with staged establishment of precursor-generating capacity during maturation (Fig. 4e). Notably, polyketide pathway genes (*PK4*; LOC115699293*, OLS*; LOC115700696*, OAC*; LOC115723437, *OAC*; LOC115723438) followed kinetics more similar to fatty acid/hexanoate-associated genes than to cannabinoid synthases. These genes were induced prior to maximal cannabinoid synthase expression, with cannabinoid synthase transcripts (*PT1*; LOC115713215, *PT7*; LOC115713205) highly expressed in late disc states. This suggests establishment of polyketide and acyl precursor capacity before full secretory maturation. However, *PT1* and *PT7* did not increase monotonically. Rather, *PT1* expression rose sharply during the transition into active secretory disc cells, peaked, and then declined in the most terminal pseudotime states while *PT7* followed a similar expression pattern then plateaued in terminal disc cells. Monoterpene synthases displayed a closely similar kinetic profile, with either a coordinated late induction followed by attenuation at the extreme end of the trajectory or a terminal disc cell expression plateau. Taken together, we suggest that terpene and cannabigerolic acid biosynthetic capacity contains both transient and sustained components across glandular trichome development.

Genes involved in isoprenoid pre-cursor flux including geranyl diphosphate (GPP) biosynthesis (*GPPSss*; LOC115725388) and an *FNR-root isozyme* (LOC115697214) exhibited broader and more sustained induction along the trajectory, consistent with progressive enhancement of isoprenoid precursor supply (Fig. 4e and Extended Data Fig. 5b)^69–71^. Their expression dynamics likely reflect increasing flux demand imposed by the simultaneous activation of both monoterpene and cannabinoid pathways, which converge on GPP-derived intermediates. Together, these data support a model in which disc cell maturation involves coordinated upscaling of isoprenoid precursor capacity to meet dual pathway demand, followed by peak activation of terminal synthases and subsequent transcriptional attenuation in fully mature secretory disc cells.

### Cell type- and stage-specific transcription factor modules coordinate metabolic and secretory programs in glandular trichomes

We next sought to define glandular trichome cell type-specific transcriptional networks by studying the TFs expressed in these cell types. We selected twelve candidates by identifying TFs with structured expression (Moran’s *I* > 0.1, *q* < 0.01) along the glandular trichome lineage trajectory, suggestive of activity at different stages of development, with emphasis on disc and neck cell enriched TFs (Supplementary Table 4a). These were four disc cell-specific (EOT1; LOC115701393, SHI6; LOC115707072, WRI1; LOC115706982, WRKY9; LOC115725761) and three neck cell enriched TFs (WRKY12; LOC115706613, MYB102; LOC115698973, and AP2-ERF; LOC115699238), together with five TFs expressed in protodermal and anthocyanin containing epidermal cells (YABBY5; LOC115724567, MYB15; LOC115704946, MADS3; LOC115704294, MYB113; LOC115708959, and MYB102; LOC115704690). We then determined their binding sites genome-wide by performing DNA affinity purification sequencing (DAP-seq)^72^. All twelve proteins were successfully expressed at the expected molecular weight (Extended Data Fig. 7a, Supplementary Table 5a). Following irreproducible discovery rate (IDR) filtering and fraction of reads in peaks (FriP) thresholding (>5% reads in IDR peaks), four high confidence datasets were retained for downstream analysis; two disc cell-enriched TFs (EOT1; LOC115701393, WRKY9; LOC115725761) and two neck cell-enriched TFs (MYB102; LOC115698973, and WRKY12; LOC115706613) (Fig. 5a, Extended Data Fig. 7b, and Supplemental Table 5b-f)^73^. We refined the list of DAP-seq-identified binding sites by cross-referencing them against the snATAC-seq data, retaining only binding sites located in accessible regions of the genome for the trichome lineage. Thereby we identified 1,707 replicable and accessible EOT1 binding sites in the trichome lineage (of 13,409 total), 1,897 replicable and accessible MYB102 peaks (of 14,519 total), 1,823 replicable and accessible WRKY9 binding sites (of 11,529 total), and 2,002 replicable and accessible WRKY12 binding sites (of 12,515 total) (Extended Data Fig. 7c).

**Fig. 5.**
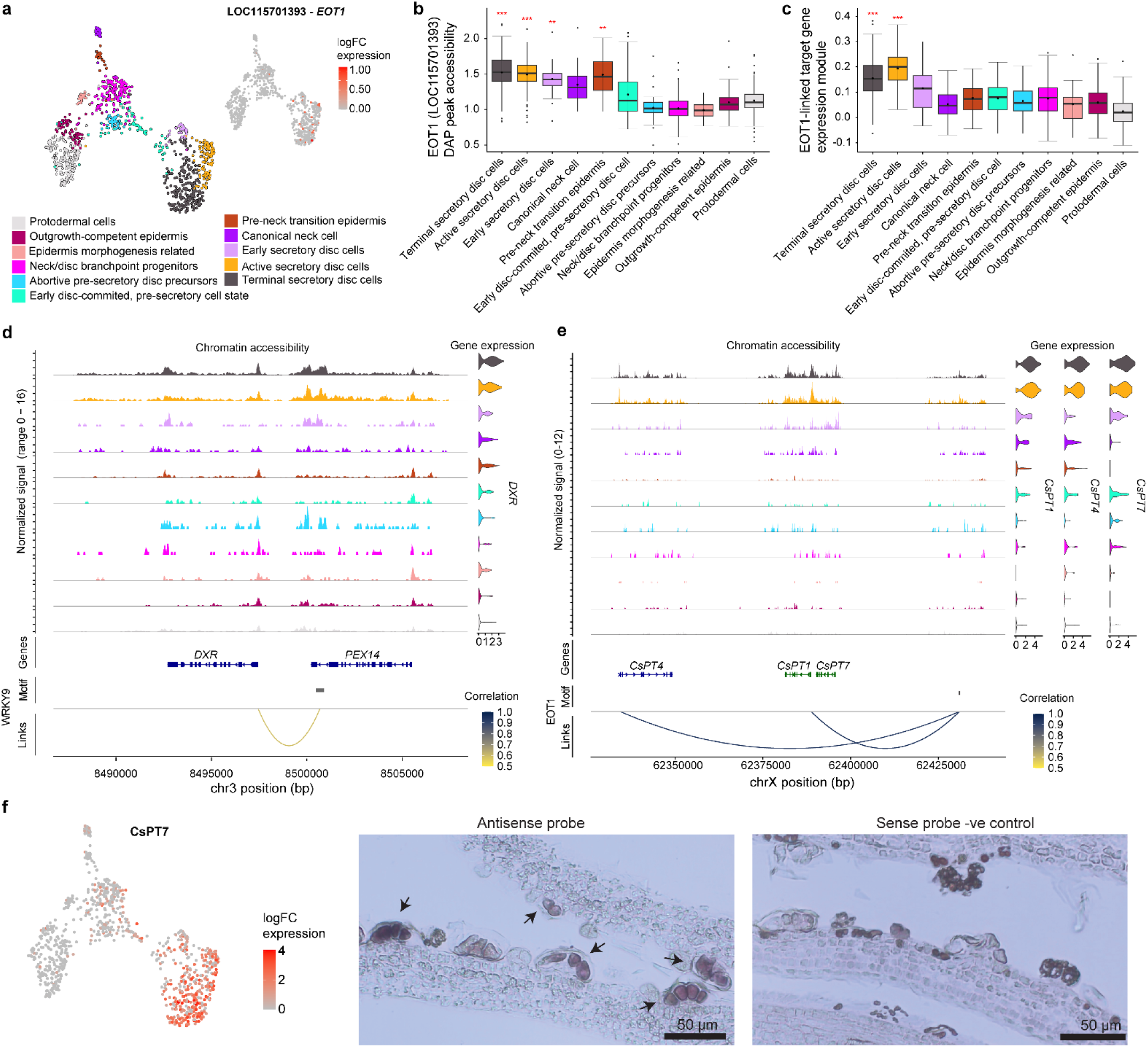
| Cell type- and stage-specific transcriptions factors coordinate metabolic and secretory programmes in glandular trichomes. **a**, UMAP of trichome-lineage cell showing *EOT1* (LOC115701393) expression, which is restricted to mature secretory disc cell states. **b**, Boxplots showing enrichment of accessible chromatin peaks containing *EOT1* binding motifs across-trichome lineage cell types. significance was assessed using two-sided Wilcoxon tests with adjusted *P* values (* < 0.05, ** < 0.01, *** < 0.0001). **c**, Boxplots showing expression of an *EOT1* motif-associated target gene module across trichome-lineage cell types. Target genes were inferred from accessible peaks containing *EOT1* motifs in disc-cell programme cells and were enriched in active and terminal secretory disc cells. significance was assessed using two-sided Wilcoxon tests with adjusted *P* values (* < 0.05, ** < 0.01, *** < 0.0001). **d**, scMultiMap peak-gene linkage highlighting a disc-cell accessible peak containing a WRKY9 (LOC115725761) binding motif associated with *DXR* expression; the linked peak lies at the 3’end of the neighbouring gene *PEX14*. **e**, scMultiMap peak-gene linkage highlighting a distal accessible peak containing and *EOT1* binding motif positively associated with *CsPT1* and *CsPT4* expression. **f**, *CsPT7* expression is restricted to mature disc-cell populations and is validated by RNA *in situ* hybridisation, which shows signal confined to mature glandular trichome disc cells.

We verified the quality of the DAP-seq data by examining motif enrichment of the accessible binding sites, then retaining the motif coordinates where the cognate motif is confidently identified in peaks for downstream analyses (Extended Data Fig. 7d,e, 8, Supplementary Table 5g-k). EOT1 DAP-seq binding sites and reciprocal accessible regions were centrally enriched with a palindromic sequence with a CTAG core motif that had degenerate flanking positions, and which had similarity with MYB3 MA1038.2 (e-value 1.8E-02) and FaEOBII motif MA1408.1 (e-value 3.9.E-01). The EOT1-associated motif contains a short TAG/CTAG-like core reminiscent of the broader, degenerate sequence space recognised by some plant MYB-family proteins sufficient to be weakly detected during motif scanning; however, EOT1 belongs to the SRS zinc-finger family rather than MYB family^74^. Consistent with EOT1 motif enrichment, SRS factors have been reported to bind a palindromic sequence with a CTAG-containing core motif, suggesting SRS-family DNA recognition may tolerate substantial motif degeneracy^75^. MYB102 binding sites were centrally enriched with a G-rich motif with a central GGTAGGT-like core, consistent with canonical MYB-binding elements and highly similar to MYB111 motif MA1036.1 (e-value 4.5E-05). WRKY9 and WRKY12 binding sites were centrally enriched with a W-box motif (TTGAC[T/C]), consistent with canonical WRKY-binding sequences and highly similar to WRKY55 (e-value 2.2.E-04) and WRKY71 motif MA1316.2 (e-value 2.7E-05), respectively (Extended Data Fig. 7d, Supplementary Table 5g-k)^22,76,77^. For each transcription factor we retained only those peaks and motif coordinates wherein the cognate motif could confidently be assigned, for downstream analyses.

We next asked if TF binding site accessibility was enriched in the cell type where the TF was expressed (Fig. 5a,b, Extended Data Fig. 9a). Accessibility of binding sites for EOT1 (disc cell TF) was significantly enriched (Wilcox two-sided adjusted *P* < 0.05) in the three secretory disc cell types (early, active and terminal), and also in pre-neck transition epidermis cells. Accessibility of binding sites for WRKY9 (disc cell TF) was significantly enriched in early, active and terminal secretory disc cells, as well as the abortive pre-secretory disc precursors and the canonical neck cells. Accessibility of binding sites for MYB102 (neck cell TF) was significantly enriched in canonical neck cells as well as active and terminal secretory disc cells. Accessibility of binding sites for WRKY12 (neck cell TF) was significantly enriched in canonical neck cells, but also in early, active and terminal secretory disc cells, and abortive pre-secretory disc precursors. These observations are broadly, but not perfectly, consistent with the expression patterns of each TF (Fig. 5a, Extended Fig. 9a,b, Supplementary Table 5l). They consequently support partially the proposed cell-specific roles of the TFs. However, they may also indicate the technical limits of DAP-seq due to the assays’ *in vitro* nature, as well as the typical property that individual genes are regulated by multiple TFs with closely spaced binding sites meaning that other factors may also influence accessibility of those locations^17,78^.

If a TF is active within a specific cell-type, we would expect expression of the TF’s target genes to be enriched within that cell-type. We examined whether this was the case for our four candidate TFs. First, target genes associated with accessible TF binding sites were identified in each cell type of the neck cell and disc cell programs (Supplemental Table 5m-u). Then the gene module score (a population-level metric of gene expression) was computed for target genes in each cell type (Fig. 5c and Extended Data Fig. 9c).To investigate each TF within the chromatin context of the cell it is expressed we modelled TF gene targets according to accessible chromatin from either the disc cell program cells for EOT1 and WRKY9 TFs or neck cell program cells for MYB102 and WRKY12. To do this we subset the disc cell program cells and recalled peaks across samples to create a unified peak list of 24,782 peaks representative of the disc cell chromatin landscape while for neck cell program cells peak calling was underpowered due to both low cell number and sparsity of snATAC data producing 3,297 accessible peaks, to improve peak calling we decided to recall peaks on the merged fragment files of the neck cell program cells which generated 6,999 representative peaks^79^. Next, we filtered previously defined EOT1 and WRKY9 accessible motifs by disc cell program peaks, and MYB102 and WRKY12 by neck cell program peaks then identified gene targets using two approaches first by retaining those expressed genes with an accessible motif in their regulatory elements or by retaining gene targets defined by scMultiMap analysis of gene-enhancer interactions in the trichome lineage (Supplemental Table 5m-u). Target-gene module expression scores across the trichome lineage broadly resembled each TFs expression patterns (Fig. 5c and Extended Data Fig. 9b,c). Expression of the target genes of the disc cell TFs was significantly enriched in terminal and active secretory disc cells (EOT1) and early secretory disc cells (WRKY9), respectively. Expression of the target genes of neck cell TF MYB102 was highest and significantly enriched in canonical neck cells, and significantly enriched with lower expression in pre-neck transition epidermis and active secretory disc cells. Expression of the target genes of neck cell TF WRKY12 was highest and significantly enriched in pre-neck transition epidermis, and significantly enriched with lower expression in canonical neck cells and active secretory disc cells.

Next, we investigated the gene targets using a combination of manual inspection and GO BP analysis to understand what processes may be coordinated by each of the four TFs in the context of the neck and disc programs. Considering first the neck cell TFs, WRKY12 target genes were suggestive of early cell entry to phenylpropanoid metabolism (*4CL*) (Supplemental Table 5m-u). Despite containing only 28 genes, the MYB102-associated neck-cell target gene module contained multiple genes linked to shikimate-derived precursor metabolism (*chorismate synthase*, *shikimate kinase*) alongside key entry points into phenylpropanoid biosynthesis (*PA*L, *4CL*), early flavonoid metabolism (*CHI*), and the lignin monomer methylation related *SAMS1*^80–83^. The recovery of multiple components spanning this metabolic continuum is unlikely to be coincidental and supports enrichment of phenylpropanoid-related processes in neck cells. Given that phenylpropanoid intermediates also feed into lignin biosynthesis these data are consistent with a broader role for neck cells in supporting aromatic compound metabolism and cell-wall associated processes (Extended Data Fig. 9 d,e). Taken together, the data suggest that WRKY12 primes early neck development and phenylpropanoid biosynthesis in the pre-neck transition epidermis while MYB102 expression demarcates mature canonical neck cell activity, producing phenylpropanoid precursors for lignin biosynthesis and barrier forming identity associated with neck cells^32^.

Gene targets of the disc cell TF WRKY9 did not enrich for any significant GO BP terms however, inspection of the target genes revealed genes involved in vesicle trafficking and membrane organization, including RAB GTPases, exocyst components, and SNARE-associated factors, together with calcium-dependent signalling modules and additionally linked to expression of the early MEP pathway gene *DXR* (Fig. 5d and Supplementary Table 5o,p)^84–87^. Together with the target gene expression patterns, these data suggest that WRKY9 may define a transitional program in which disc cells are primed for secretion through establishment of trafficking capacity, membrane polarity, and signalling competence, preceding maximal secretory output.

Gene targets of the disc cell TF EOT1 were enriched for terpenoid, isoprenoid, and cannabinoid biosynthetic processes (Extended Data Fig. 9e and Supplemental Table 5v). Notably, among the strongest expression-accessibility correlations observed (Pearson r ≈ 0.8 - 0.95, adjusted *P* < 0.05) were associations between EOT1 binding sites were linked to CBGAS genes *PT1* and *PT4*, and cannflavin-related prenyltransferase *PT3* (Fig. 5e and Supplementary Table 5n)^88,89^. *PT1, PT3,* and *PT4* form part of a large cluster (∼342 Kb) of six CBGA-like prenyltransferase genes. We examined the gene cluster closely, determining that the expression of five genes were linked to a single accessible EOT1 site (chrX-62430871-62431241) and that all 6 members of this cluster are co-expressed in the disc cell program (Fig.5e and Supplementary Table 4a,b). We sought to validate the spatial restriction of this co-expressed cluster of genes to disc cells, so selected one gene (*PT7,* LOC115713205) from the cluster and analysed its expression pattern by RNA *in situ* hybridization. This confirmed expression was restricted to mature disc cells (Fig. 5f)). These findings support a model in which EOT1 activity is coupled to activation of a prenyltransferase module in mature disc cells, directly linking TF occupancy to cannabinoid pathway deployment during terminal secretory differentiation of this cell type.

In parallel, we also observed that EOT1 target genes encoding downstream metabolic enzymes, including *NES*, *myrcene synthase*, and *licodione synthase* indicating association with metabolic output rather than pathway initiation alone^90^ (Supplemental Table 5n). We noted the presence of the plant oil biosynthesis master regulator *WRI1*^62^. This was accompanied by genes supporting acetyl-CoA / terpenoid precursor supply, including *ACAT, FBA3, IDH,* and *MPC4*, consistent with strong biosynthetic commitment^91–93^. Notably, EOT1 targets also included transporters and membrane-associated factors such as *ABCG11*, *PDR1*, and *SWEET16-like*, together supporting a model in which EOT1 drives high sink strength and secretory capacity in active and terminal disc cells^94–96^. In contrast to the earlier WRKY9-associated state, which was linked to precursor-setting functions including *DXR*, the EOT1 program is associated with terminal metabolic biosynthesis capacity exemplified by *myrcene synthase*, *PT4,* and *PT1*, and with the transport and metabolic infrastructure required to sustain secretion (Supplementary Table 5m-p).

### MeJA signalling remodels disc cell specialised metabolism and reallocates resources to extracellular defence

To examine transcriptional and chromatin responses associated with early defence elicitation, we compared gene expression between single-cell transcriptomes from control and 2-h MeJA (0.02%, ∼1 mM) treated *C. sativa* inflorescences. Two gene modules were identified by differential gene expression analysis of pseudobulked control and treatment data, comprising transcripts either induced or suppressed following treatment, with 70 genes significantly upregulated and 53 genes downregulated (Supplementary Table 6a). We examined the cell type specificity of differential expression by projection of the modules across inflorescence cell identities. This revealed contrasting spatial patterns. Genes induced following methyl jasmonate treatment showed a broadly uniform enrichment across diverse floral cell types, consistent with activation of a widespread defence program (Fig. 6a). In contrast, genes downregulated after treatment exhibited strong enrichment within the glandular trichome disc cell identity (Fig. 6b). We then mapped the induced gene module onto the trichome-associated epidermal atlas which revealed comparatively broad enrichment across cell types except for pre-neck transition epidermis and abortive pre-secretory disc cells (Fig. 6c). Mapping the repressed module onto the trichome-associated epidermal atlas further revealed that mature secretory disc cells- particularly active and terminal secretory disc cells- disproportionately contribute transcripts suppressed following MeJA treatment (Fig. 6d).

**Fig. 6.**
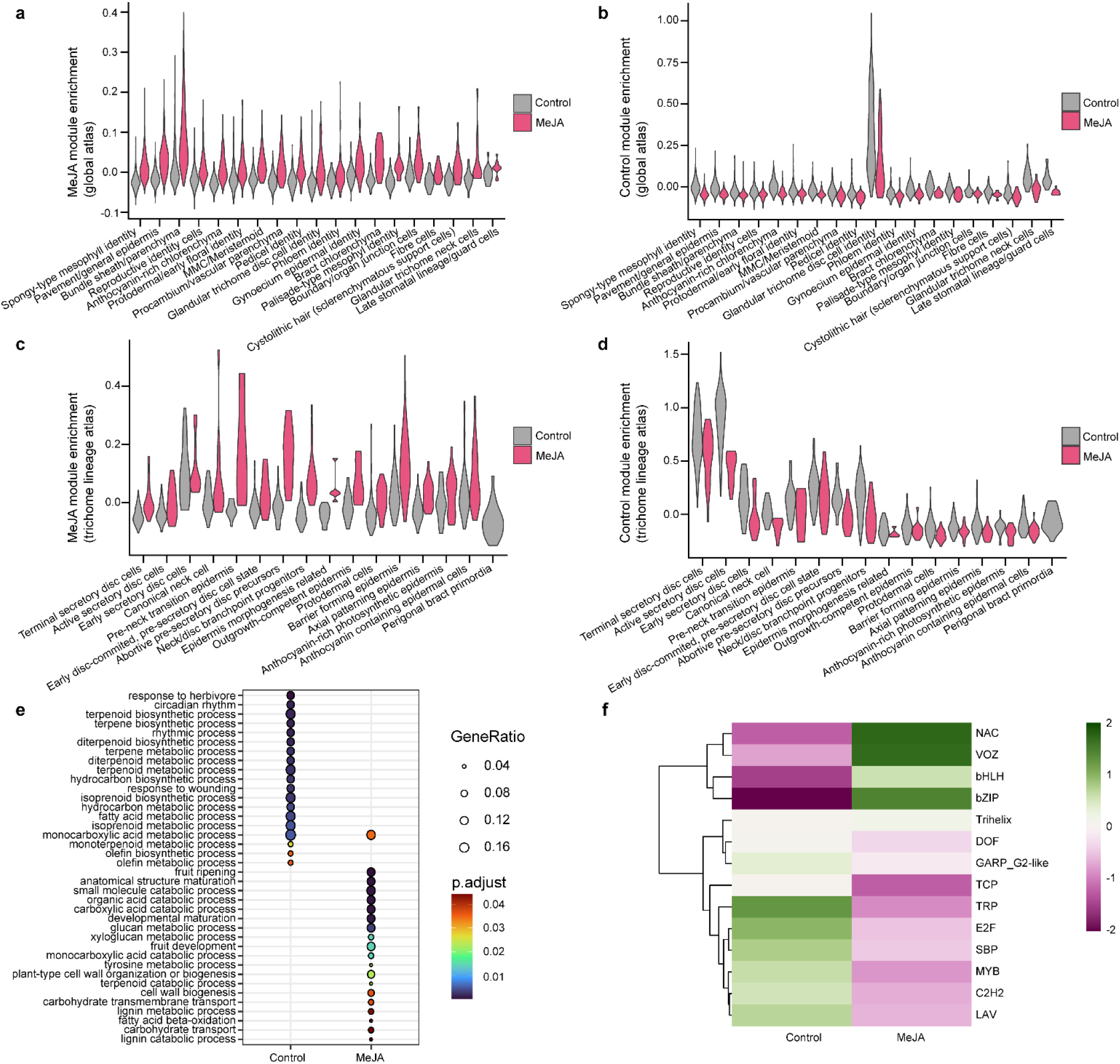
| Acute jasmonate signalling induces a broad extracellular defence programme while selectively suppressing disc cell terpenoid specialised metabolism. **a**, Violin plots showing module score for genes upregulated in methyl jasmonate (MeJA)-treated inflorescence pseudobulk samples across cell types of the inflorescence atlas. MeJA-upregulated genes are broadly distributed across cell types. **b**, Violin plots showing module scores for genes upregulated in control inflorescence pseudobulk samples across cell types of the inflorescence atlas. Control-upregulated genes are enriched in glandular trichome disc-cell populations. **c**, Violin plots showing module scores for MeJA-upregulated pseudobulk genes across cells in the trichome-associated epidermal atlas. MeJA-upregulated genes are broadly distributed across epidermal states. **d**, Violin plots showing module scores for control-upregulated pseudobulk genes across cells in the trichome-associated epidermal atlas. Control-upregulated genes are enriched in active and terminal secretory disc cells. **e**, Gene ontology biological process enrichment of control-upregulated and MeJA-upregulated inflorescence pseudobulk genes (adjusted *P* < 0.05). Control-upregulated genes are enriched for specialised metabolic and trichome-associated processes, whereas MeJA-upregulated genes are enriched for extracellular, cell wall-related and resource reallocation processes. **f**, Heatmap of chromVAR-derived transcriptiona factor motif family enrichment (JASPAR 2024) across control and MeJA-treated samples, shown as Z-score-scaled accessibility.

The functions of genes differentially regulated by MeJA were assessed using GO analysis. Upregulated genes were enriched for extracellular and cell-wall-associated processes, including cell wall polysaccharide metabolism (for example *laccase-7*; LOC115696962, *polygalacturonase*; LOC115708768) together with carbohydrate remobilization pathways including β-amylase 3 (LOC115716102)^97–99^ (Fig. 6e and Supplemental Table 6b). They were also enriched for fruit ripening and fruit development terms, processes that MeJA has been linked to in multiple species and suggesting the same may be true for *C. sativa*^100,101^. By contrast, downregulated genes were enriched for specialised metabolic pathways including terpenoid and flavonoid biosynthesis, as well as genes associated with hexanoate (*HCS/AAE1*; LOC115709751, *OLS*; LOC115700696), cannflavin (*PT3*; LOC115713148), and monoterpene (*CsTPS6FN*; LOC115716604, *CsTPS6FN;* LOC115716405) metabolism^5,24,89^ (Fig. 6e and Supplemental Table 6b). Counterintuitively, we observed enrichment of a herbivory response GO term, comprised of two sesquiterpene and two monoterpene synthase genes. Interestingly, despite strong enrichment of disc cell terminal specialised metabolism genes amongst the downregulated genes, including the *PT3* cannflavin member of the CBGAS gene cluster, we did not observe any effect of MeJA treatment on canonical terminal cannabinoid biosynthetic pathway genes. Indeed, canonical CBGA synthases *PT1* and *PT4* were previously shown to lack jasmonate responsive elements in their regulatory elements^88^. Together these data suggest *C. sativa* responds to acute MeJA by rewiring disc cell specialised metabolism such that cannabinoid synthase transcription remains unperturbed while monoterpene, sesquiterpene, and flavonoid metabolism is suppressed.

Next, we asked if changes in chromatin accessibility supported the observed transcriptomic changes following MeJA treatment. To this end, ATAC-seq peaks from each cell type of the trichome associated epidermal cells that were then analysed under control and treatment conditions. Coordinated changes in TF motif accessibility were observed following MeJA treatment by applying ChromVar analysis (Fig. 6f)^36^. Motifs associated with stress-responsive TF families, including NAC, bZIP, and VOZ, displayed increased accessibility in MeJA-treated cells, consistent with activation of defence associated transcriptional programs^102–104^. In contrast, motifs corresponding to developmental and metabolic regulators, including TCP, MYB, SBP family TFs, showed reduced accessibility^105–107^. Notably, several of these families have been implicated in the regulation of specialised metabolism and epidermal differentiation, suggesting that jasmonate signalling represses transcriptional networks associated with secretory activity while activating stress-responsive regulatory programs. Together, these results suggest that jasmonate signalling may indirectly enhance cannabinoid specialised metabolism by reducing competition for isoprenoid precursors through the attenuation of monoterpene synthases.

## Discussion

Our study provides a cell-resolved model for the ontogeny of *C. sativa* glandular trichomes. Rather than representing distinct glandular trichome classes, bulbous, pre-stalked, and capitate-stalked morphotypes are more consistent with sequential stages along a single developmental lineage originating from the protoderm and culminating in terminal capitate-stalked glandular trichomes. This framework reconciles previous morphological observations with a developmental model and provides an explanation for how a spatially uniform protoderm gives rise to large pre-stalked glandular structures. In particular, our data support the proposal that pre-stalked trichome are intermediates preceding capitate-stalked trichome differentiation and further suggest that bulbous trichomes represent an earlier transitional stage within the same lineage^18,26,27^.

Within this framework, we identify and validate the glandular trichome neck cell as a transcriptionally distinct cell type arising from an early bifurcation in the protoderm-to-disc-cell trajectory. Although neck cells have been inferred previously from histological observations, our data place them explicitly within the glandular trichome developmental lineage and suggest they are specified before terminal secretory differentiation^32,108^. The enrichment of suberin- and lignin-associated regulatory programs further suggests that neck cells contribute to the structural and physiological segregation of the gland head from underlying tissues^32,109,110^. This raises the possibility that neck cells act not only as a physical boundary but also as a specialised diffusion barrier that supports the compartmentalisation of secretory metabolism within the mature gland head.

Our analyses suggest that glandular trichome differentiation is not governed by a single shared regulatory program, but by parallel state-specific TF modules operating within the neck and disc lineages. Together, these analyses support a model in which glandular trichome differentiation is coordinated by cell type- and stage-specific TF modules that partition metabolic, structural, and secretory functions between neck and disc cell states. In disc cells, the earlier WRKY9-associated program is linked to signalling, membrane trafficking and acquisition of secretory competence, whereas the EOT1-associated program is coupled more strongly to downstream specialised metabolism, transport and high secretory throughput, consistent with active metabolite production and export. In this context EOT1 does not simply activate terminal specialised metabolic enzymes, but instead orchestrates the metabolic, redox, and secretory capacity required to sustain high-flux cannabinoid and terpenoid biosynthesis in secretory disc cells. By contrast, the neck-cell regulators define a distinct but complimentary axis: WRKY12-asssociated targets are enriched in the pre-neck transition epidermal state and emphasise receptor mediated signalling, redox buffering, and phenylpropanoid entry-point enzymes, while MYB102-associated targets define the mature canonical neck cell state and are enriched for shikimate and phenylpropanoid metabolism, including lignin-associated pathways consistent with the formation of a lignified barrier domain. Thus, rather than reflecting a single uniform glandular trichome program, these data indicate that neck and disc cells are established through parallel regulatory modules, with disc-cell TFs preferentially reinforcing metabolic output and secretion, and neck-cell TFs reinforcing signalling transitions and lignin-associated barrier formation required to support glandular trichome function.

Considering glandular trichomes are major defensive structures, understanding how defence signalling reshapes their regulatory state is an important outstanding question. At single-cell resolution, acute MeJA elicitation revealed extensive transcriptional and chromatin rewiring in mature disc cells that was accompanied by repression of specific specialised metabolic programs and activation of broader defence- and cell wall-associated responses. Cannabinoids are well documented for their potent role in anti-herbivory and we speculated this specialised metabolism transcriptional shift could indirectly increase cannabinoid biosynthesis due to reduced competition for isoprenoid precursors. This model is supported by previous observations of increased cannabinoid content and unperturbed expression of cannabinoid biosynthetic genes following MeJA treatment in *C. sativa*^24,111–114^. Together, our results establish a cell-resolved framework for glandular trichome development, secretory cell specialisation and defence-associated regulatory plasticity in *C. sativa.* Beyond refining the developmental and cell-biological model of *C. sativa* glandular trichomes, these findings provide a foundation for engineering glandular trichomes as platforms for the high-yield production and storage of valuable specialised metabolites.

## Methods

### Micro-computed tomography (Micro-CT)

Clones of C. sativa (MW-615) were grown in 4.5-L pots containing Rotterdam 70:30 coco-perlite soil media in a controlled environment room (CER; 12 h of light at 28 °C / 12 h of dark at 22°C with 55 % humidity and an average light intensity of 250 µmol m−2 s−1). Nutrient feeding regimes were as described by Welling et al^24^. Subapical floral tissues (bract, ovary, stigma) from 4-, 5-, or 9-week-old flowering plants (n = 2 biological plant replicates) were snap-frozen in liquid nitrogen, thawed immediately before imaging, and mounted at 24 °C (room temperature) in sealed polypropylene tubes using a water-based adhesive. X-ray phase-contrast micro-CT imaging was performed at the Micro-Computed Tomography (MCT) beamline (Hutch B) of the Australian Synchrotron (ANSTO)2. Samples were imaged using a monochromatic 17 keV beam and a 4.533 × objective on the white detector (PCO.edge 5.5) at a 160 mm sample-to-detector distance, yielding a 3.7 × 3.1 mm field of view and an effective pixel size of 1.434 µm. Scans were acquired in continuous fly-scan mode over 183°, using 1831 projections with 0.025 s exposure. Projection data were reconstructed with the Australian Synchrotron MCT pipeline using TIE-HOM phase retrieval, as described by Arhatari et al.^115^, producing isotropic voxels of 1.42 µm. Reconstructed volumes were visualized in ORS Dragonfly Pro v2022.2 using the MCT import script (https://github.com/AustralianSynchrotron/dragonfly_mct_import).

### Plant growth conditions

Cuttings were taken from female genotype (MW6-15) industrial hemp line (Accession #6) mother plants. Cuttings were made using a sterile blade and at a 45-degree angle immediately below the 3^rd^ node from the apical meristem. The cuttings were partially pruned – removing leaf tips and large leaves, taking care not to remove all leaves. The cuttings were then dipped in 3.0 g/L indole butyric acid rooting gel hormone (growth technology © Clonex purple) and allowed to sit for 30 seconds. Cuttings were individually transferred to Grodan Rockwool cubes (36mm x 36mm x 40mm) previously saturated with half strength CANNA veg fertiliser (20 mL A + 20 mL B/ 10 L water) and then transferred to propagator humidity domes (4 clones to one propagator) with the vent fully closed. Clones were monitored daily – half strength CANNA veg fertiliser (20 mL A + 20 mL B/ 10 L water) was added to the base of the propagator when required and vents were opened in small daily increments until completely opened and grown under vegetative lighting conditions of 18 hours on 6 hours off using Philips Master TL-D Super 80 low-pressure mercury discharge lamps. Clones were kept in propagators until established roots could be observed (approximately 2 weeks) at the base of the rockwool cubes. Potting mix containing 1:1:1 peat moss, perlite, and vermiculite supplemented with 1 g/L dolomite was prepared and divided into 1 L pots. Pots were then saturated with full strength CANNA veg fertiliser (40 mL A + 40 mL B/ 10 L water). Clones were then transferred to appropriately labelled 1 L pots and allowed to grow under vegetative lighting conditions for 4 weeks. Plants were monitored daily and watered regularly using full strength CANNA veg during the vegetative growth stage. Following 4 weeks of vegetative growth plants were re-potted into appropriately labelled 10 L pots using the above potting mix recipe. Potting mix was saturated with full strength CANNA flora (40 mL A + 40 mL B/ 10 L water) prior to re-potting of the clones. The lighting cycle was then varied to 12 hours on and twelve hours off to induce flowering. Clones were grown under flowering conditions for 6 weeks and watered daily with full strength CANNA flora.

### Methyl jasmonate (MeJA) treatment

At 6 weeks post floral induction (PFI) plants were treated with either 1mM MeJA dissolved in RO water supplemented with 0.8% (v/v) EtOH and 0.1% (v/v) Tween 20 or control RO supplemented with 0.8% (v/v) EtOH and 0.1% (v/v) Tween 20 conditions^24^. Two biological replicates were performed for both treatment and control. Treatment and control were applied as a whole plant foliar spray where each biological replicate received 25 ml. Treatment and control were applied in separate growing chambers to prevent cross contamination. Inflorescence tissue was sampled for downstream nuclei isolation following 2 hours of treatment and control application.

### Nuclei isolation

To isolate high-quality plant nuclei compatible with the downstream 10X Genomics snMultiome workflow we modified a tomato green tissue nuclei isolation protocol^116^. A step by step procedure for the modified protocol was then made available^117^. The following buffers and Percoll® (Sigma-Aldrich P4937) solutions were prepared fresh on the day of experiment and RNase protector inhibitor (Roche 03335399001) was added to each buffer immediately prior to use. All buffers and Percoll® solutions were chilled in ice prior to use.

Nuclei wash buffer (NWB) consisted of 10 mM Tris-HCl (pH 6.9), 10 mM NaCl, 3 mM MgCl_2_, 5 mM CaCl_2_, 1% BSA (Sigma-Aldrich A9647), 0.1% Tween-20 (Bio Rad 1610781), 1 mM DTT (Thermo Fisher Scientific A39255), 1 U μl^−1^ Protector RNase inhibitor; 30% Percoll® solution consisted of 70% nuclei wash buffer, 30% Percoll® (v/v), and 1 U μl^−1^ Protector RNase inhibitor; 70% Percoll® solution consisted of 30% nuclei wash buffer, 70% Percoll® (v/v), and 1 U μl^−1^ Protector RNase inhibitor. Nuclei lysis buffer (NLB) consisted of 10 mM Tris-HCl (pH 6.9), 10 mM NaCl, 3 mM MgCl_2_, 5 mM CaCl_2_, 1% BSA, 0.1% Tween-20, 0.2% Triton X-100 (Sigma-Aldrich X-100), 1 mM DTT, 1 U μl^−1^ Protector RNase inhibitor. Diluted 10x Genomics nuclei buffer consisted of 1X 10x Genomics 20X concentrated Nuclei buffer (10x Genomics PN-2000207), 1 mM DTT, and 1 U μl^−1^ Protector RNase inhibitor. All nuclei isolation steps were carried out on ice. For each sample, 200 mg of fresh *C. sativa* inflorescence was chopped for 2 min using a razor blade in a petri dish with 400 μl of NWB, then a further 350 μl of NWB was added and the cell lysate was allowed to rest on ice with gentle agitation for 2 min. The petri dish was tilted at a 45° angle, and cell lysates were collected using a p1000 pipette with wide bore tips, then transferred into pre-wetted 20 μm mesh size mini pluristrainers (pluriSelect 43-10020-50), and the filtrates were collected in 1.5 ml DNA LoBind® Eppendorf tubes (Eppendorf 022431021). Filtrates were then centrifuged at 50g for 3 min at 4°C using a swinging bucket rotor to pellet large debris. 500 μl of each sample supernatant was carefully aspirated using a wide bore pipette tip, so as not to disturb the debris pellet, then filtered into a new 1.5 ml LoBind® Eppendorf tube using a pre-wetted 20 μm mesh size mini pluristrainer. Density gradients were then established by slowly underlaying 200 μl of 30% Percoll® solution followed by 200 μl of 70% Percoll® solution beneath the 30% Percoll® layer. Density gradients were then centrifuged at 2500g for 15 min at 4°C. 200 μl of the green band formed between 30% and 70% Percoll® layers, containing high quality nuclei and minor chloroplast contamination, were aspirated and transferred to a new 1.5 ml LoBind® tube. To remove chloroplast contamination and permeabilise the nuclear membrane for downstream transposition 1 ml of NLB was added to each tube and the suspension was allowed to rest on ice for 5 min with gentle inversion every 1 min. The lysate was then filtered into a 1.5 ml LoBind® tube using a pre-wetted 10 μm mesh size mini pluristrainer (pluriSelect 43-10010-50), and then centrifuged at 500g for 5 min at 4°C. The supernatants were then removed, and the nuclei pellets were resuspended in 1 ml of NWB by gentle pipetting 5X, then centrifuged at 500g for 5 min at 4°C, wash was repeated 2 more times. The pellet was then resuspended in 100 μl of diluted 10x Genomics nuclei buffer and centrifuged at 500g for 5 min at 4°C. The supernatants were removed and the nuclei pellet was resuspended in 20 μl of diluted 10x Genomics nuclei buffer and filtered into a 1.5 mL LoBind® tube using a pre-wetted 10 μm mesh size mini pluristrainer then placed on ice. A 5 μl aliquot, of each nuclei preparation, was transferred to a new 1.5 ml LoBind® tube for downstream tagmentation.

Nuclei quality and quantity were assessed by taking 10 μl of the remaining nuclei preparation followed by the addition of 0.5 μl 50 μg ml^-1^ Hoescht 33342 (Thermo Fischer Scientific H3570), then assessing the nuclei quality by fluorescence microscopy and estimating counts using a haemocytometer.

### Single-nucleus multiome library preparation

Same cell single-nucleus assay for transposase accessible chromatin and gene expression libraries from control and MeJA treated *C. sativa* inflorescence tissues were prepared as follows. After estimating nuclei counts, unstained aliquots were diluted using dilute 10x nuclei buffer to obtain ∼ 16,000 nuclei in 5 μl for each sample. Nuclei were then Tagmented, partitioned, and barcoded using a Chromium X series controller (10x Genomics) and a Chromium Next GEM Single Cell Multiome ATAC + Gene Expression Reagent kit (10x Genomics) following the manufacturers protocol. Single-nucleus ATAC and gene expression sequencing libraries were then generated according to the manufacturers protocol. Sequencing library quality was inspected on a 4150 Agilent Tapestation system using high sensitivity DNA 5000 screen tapes (Agilent). Sequencing for snATAC libraries was performed on an Illumina NovaSeq 6000 S1 flow cell using 2x 50 bp paired-end sequencing, targeting 500,000,000 read pairs per sample. Sequencing for snRNA libraries was carried out on a NovaSeq X 10B flow cell using 2x 150 bp paired-end sequencing, targeting 300,000,000 read pairs per sample.

### Single-nucleus ATAC + RNA-seq analysis

CellRanger ARC (v. 2.0.2) was used to generate raw UMI matrices for each sample separately, mapped to cs10 reference genome (GCF_900626175.2) supplemented with the *C. sativa* Yunma 7 chloroplast genome (MW013540)^118–120^.

### Background RNA contamination correction

We applied CellBender (v. 0.3.2) on the raw feature barcode matrix with the option exclude-feature-types Peaks to remove both empty droplets and background RNA contamination from nuclei containing droplets. Filtered feature barcode matrix files were then loaded into Seurat (v.5.4.0)^121,122^.

### Generation of Seurat and Signac objects

Single-nucleus RNA and chromatin accessibility data were processed using Seurat (v.5.4.0) and Signac (v1.16)^79,122^. Using the Signac CreateChromatinAssay function chromatin assays were added to each sample object using the associated fragment files. Sample ID and treatment ID metadata were appended to each Seurat objects metadata.

### ATAC peak calling

Peaks were called according to the ATAC-seq peak fragment files using the MACS2 callpeak function using the following options -t -g 736579359 -f BED --nomodel --extsize 200 --shift -100 for each individual sample, then sample peak lists were combined and filtered for peaks >20 bp and < 1000 bp. Samples were then merged into a single Seurat object using the Seurat merge function and a new chromatin assay was generated using the combined MACS2 peak list^123^. Quality control was performed at the nucleus level using a combination of RNA-based, ATAC-based, and organellar contamination metrics, with thresholds defined empirically from metric distributions and applied consistently across all samples.

### Quality control filtering of cells

RNA-based filtering – Nuclei were required to meet minimum RNA complexity and abundance criteria nFeature > 200, nCount > 200. To exclude potential multiplets and high-content outliers, upper bounds were applied using the 99.5^th^ percentile of each distribution.

Nuclei with elevated ambient mitochondrial transcripts >10% were removed.

ATAC-based filtering – Chromatin accessibility quality was assessed using transcription start site (TSS) enrichment and nucleosome phasing metrics. Nuclei were retained if they met the following criteria TSS enrichment >1 and nucleosome signal <2. Upper bounds were applied to remove potential multiplets using the 99.5^th^ percentile of each distribution nCount_ATAC and nFeature_ATAC. Lower ATAC count thresholds were deliberately relaxed, as nuclei with lower fragment counts but strong TSS enrichment and low nucleosome signal were retained, indicating biologically valid chromatin accessibility profiles.

Plastid contamination-aware filtering was performed to explicitly account for plastid-reads. To do so the *C. sativa* Yunma 7 chloroplast genome (MW013540) was appended to the cs10 (GCF_900626175.2) reference genome and plastid read fractions were quantified per nucleus. Rather than applying a single hard plastid threshold, a two-step plastid/TSS-aware filtering strategy was used. Firstly, nuclei with evidence of extreme plastid transcript contamination >20% were removed^124–126^. Quality gating was then performed on nuclei with moderate plastid contamination (10-20%) where retention was conditional on chromatin quality: nuclei were retained only if TSS enrichment >2 and nucleosome signal <2. This approach ensured that nuclei with plastid reads arising from technical contamination rather than chromatin accessibility were excluded, while preserving nuclei exhibiting strong promoter accessibility and low nucleosome signal.

### Data normalisation and clustering

Chromatin accessibility matrices were normalised using term frequency-inverse document frequency (TF-IDF) followed by latent semantic indexing (LSI) as implemented in Signac. Gene expression matrices were normalised using SCTransform v2 followed by RunPCA (with 2,000 variable genes and 50 pcs)^127^. We then used the immunogenomics Local Inverse Simpson Index (iLISI) for scRNAseq data package (v. 1.0) to test median neighbourhood mixing across samples^29^. To mitigate residual sample-associated variation in RNA embeddings across four samples, we applied Harmony (v. 1.2.4) integration prior to multimodal analysis^29^. This increased iLISI from 1.72 to 2.46, indicating improved cross-sample alignment. Harmony-corrected RNA embeddings were subsequently combined with ATAC LSI components 2-30 for weighted nearest-neighbour (WNN) analysis using the FindMultiModalNeighbours function, followed by UMAP dimensionality reduction using RunUMAP^128,129^. We next used clustree (v. 0.5.1) and the FindClusters function to determine the optimal cluster resolution (0.4)^130^. Next, marker genes were identified using the FindAllMarkers (with logfc.threshold 0.1; min.pct 0.1; only.pos, T) and filtered to those with adjusted p values < 0.05, and a small cluster (218 nuclei; ∼1.8%) enriched for plastid, mitochondrial, and ribosomal protein transcripts was identified as low-quality nuclei and excluded from downstream analyses. Preprocessing steps including normalisation, dimensionality reduction, and WNN analysis, were then repeated on the newly subset data. The identity of each cluster in the global atlas was manually annotated using cell type marker genes from the literature (Supplementary Table 1e) in combination with GO enrichment analysis of the clusters^131^.

Clusters constituting the trichome lineage protodermal/early floral epidermal identity, glandular trichome disc cell identity, and glandular trichome neck cell identity, were subset into a new object using the Seurat subset function. Chromatin accessibility matrices were normalised using term frequency-inverse document frequency (TF-IDF) followed by latent semantic indexing (LSI) as implemented in Signac. Normalisation was applied to the RNA counts of the subset object using SCTransform v2 followed by RunPCA (with 2,000 variable genes and 50 pcs). To mitigate residual sample-associated variation in RNA embeddings across four samples, we applied Harmony (v. 1.2.4) integration. This increased iLISI from 2.28 to 3.06, indicating improved cross-sample alignment. To resolve finer transcriptional cell states within the trichome lineage used RunUMAP on the Harmony corrected RNA embeddings only. We next used clustree (v. 0.5.1) and the FindClusters function to determine the optimal cluster resolution (0.4). Marker genes were identified using the FindAllMarkers (with logfc.threshold 0.25; min.pct 0.2; only.pos, T) and filtered to those with adjusted p values < 0.05. The identity of each cluster in the Trichomes lineage was manually annotated using multimodal cell type markers in combination with GO enrichment analysis of the clusters.

### Monocle3 trajectory inference of the trichome lineage

Trajectory analysis was carried out using Monocle 3 (v. 1.4.26)^132^. Nuclei from the trichome lineage Seurat object were converted to a Monocle3 cell_data_set using SeuratWrappers (v. 0.4.0) as.cell_data_set function. Harmony-corrected RNA embeddings were transferred and used as the dimensional space for trajectory inference. A principal graph was learned and cells were ordered in pseudotime with protodermal cells specified as the root population^133–135^. Expression of selected genes was visualised along pseudotime using plot_genes_in_pseudotime in Monocle3. To identify transcriptional programs associated with progression along the trichome trajectory, we tested each gene for spatial autocorrelation along the learned principal graph using Monocle3 graph_test. Genes with significant pseudotime-associated structure were retained using q < 0.01 and Moran’s *I >* 0.10, then clustered into co-varying gene modules with find_gene_modules. Module activity was summarised across cells and annotated cell states, and the resulting gene sets were independently projected onto the trichome lineage atlas Seurat object using AddModuleScore. Cell states with elevated scores for a given module were considered to co-express that program, enabling modules to be assigned to discrete phases of trichome lineage progression.

### Pseudobulk differential gene expression analysis of control and treatment

Treatment response differential gene expression analysis was carried out by pseudobulking single-nucleus transcriptomes using the muscat R-package (v. 1.24.0)^136^. For each biological replicate, gene counts were summed across all cells in the inflorescence atlas to generate sample-level pseudobulk expression matrices. Differential gene expression analysis between control and MeJA- treated sample was performed using edgeR (v. 4.8.2)^137^. Library sizes were normalised using the trimmed mean of M-values (TMM) method, and genes with low expression across samples were excluded using sample level filtering implemented in muscat. Differentially expressed genes with BH adjusted p values < 0.05 were retained. Gene modules were then generated from genes significantly upregulated or downregulated in MeJA-treated samples and module scores were calculated for each cell using the AddModuleScore function in Seurat, module scores were then projected onto the trichome lineage atlas to visualise the distribution of treatment-responsive transcriptional programs across cell identities.

### chromVAR differential motif accessibility analysis

Transcription factor motif accessibility was quantified using chromVAR (v. 1.32.0)^36^. Motif deviation Z-scores were calculated for each motif across single nuclei using the chromVAR assay. Motif accessibility was evaluated separately across three peak sets: the global peak set derived from the full inflorescence atlas, re-called peaks from the trichome lineage atlas, and peaks re-called from the mature disc cells. To compare transcription factor accessibility between treatments, motif deviation scores were averaged across cells belonging to each condition. Motifs were manually annotated to transcription factor families based on motif annotations from the JASPAR 2024 database^138^. Family-level accessibility scores were calculated by averaging Z-scores across motifs belonging to the same transcription factor family. Heatmaps displaying transcription factor family accessibility across conditions were generated using the pheatmap (v. 1.0.13) R-package

### RNA *in situ* hybridization

Inflorescence tissue was harvested from *C. sativa* plants 5 weeks PFI and fixed by vacuum infiltration at 200 mbar in 4% (w/v) paraformaldehyde (Sigma Aldrich 158127) and 4% (v/v) DMSO (Sigma Aldric 34869), vacuum was released after 10 min and reapplied for an additional 10 min. Samples were stored overnight at 4°C in fresh fixative. The following day, the fixed tissues were dehydrated using a Leica TP1020 Semi-Enclosed Benchtop Tissue Processor (Leica Biosystems) at room temperature according to Liew^37,139^. Tissue was then added to molten (65°C) Surgipath Paraplast Paraffin (Leica Biosystems) twice for 2 hours each. Paraplast tissue blocks were then prepared using the Leica EG1150 H Heated Paraffin Embedding Module with the added Leica EG1150 C Cold Plate for Modular Tissue Embedding system (Leica Biosystems). Embedded tissues were sectioned at 8 µm thickness and *in situ* hybridization was carried out according to a modified protocol by Jackson^140^: 55°C hybridization temperature and 1x saline-sodium citrate (SSC) washes. Transcripts of interest were PCR amplified using primers (Supplementary Table 3) and cloned into pGEM-T Easy vector (Promega A1360). Digoxigenin (DIG)-labelled antisense and sense RNA probes were transcribed, using the DIG RNA labelling Kit (SP6/T7) (Sigma Aldrich 11175025910), from the T7 or SP6 promoter of the pGEM-T Easy vector according to the manufacturers protocol. Hybridization results were observed and imaged using a Zeiss Axio Observer A1 microscope (Carl Zeiss).

### DNA affinity purification sequencing (DAP-seq)

The DAP-seq experiment was performed using reagents as previously described, but with modifications to optimize sensitivity^72,141^. A step by step procedure for the modified protocol was then made publicly available^142^. Genomic DNA was extracted from *C. sativa* inflorescence tissue 6 weeks PFI to capture gDNA with inflorescence methylation context, using the DNeasy Plant Mini Kit (Qiagen 69106) following the manufacturers protocol, gDNA was then sheered using a Diagenode Bioruptor® UCD-200 (Diagenode) to an average fragment size of 200 bp with the following settings; water temperature 4°C, high setting, and 30 s on/off intervals over 20 min. Sheered DNA was then used as input, for library generation, using the NEBNext Ultra II DNA library prep kit (New England Biolabs inc. E7645S) following the manufacturers guidelines.

The coding sequences of *C. sativa* transcription factors (Supplementary table 5) derived from the cs10 reference genome (GCF_900626175.2) were synthesised with flanking 5’ SacI and 3’ XhoI restriction sites by Twist Bioscience (South San Francisco, CA, USA), and were cloned into a modified pIX-HALO vector. The original pIX-HALO vector sequence (signal.salk.edu) was virtually modified using Geneious Prime® (v. 2025.0.3), first the vector was linearised by removing the Gateway recombination cassette (attR1 – ccdB – attR2) and then a flexible (Gly)_4_Ser linker sequence, SacI restriction site, TAG stop codon, followed by a XhoI restriction was added to 3’ end of the Halo-tag sequence^143,144^. Then 5’ His_6_ tag sequence was removed and replaced with a XhoI restriction site. Lastly, to avoid conflicts due to long homopolymers during synthetic DNA synthesis we modified the artificial 30 bp poly A tail by introducing a G nucleotide every 12 A bps, where 11-12 consecutive A nucleotides are required for efficient binding of poly A binding protein thereby allowing efficient downstream transcription and translation while simultaneously disrupting difficult to synthesise homopolymers^145,146^. The linear sequence was then synthesised by Twist Bioscience (South San Francisco, CA, USA) and circularised by XhoI (New England Biolabs inc. R0146S) digestion of 5’ and 3’ flanks and subsequent ligation using T4 DNA ligase (New England Biolabs inc. M0202S) (Supplementary Table 5a).

A total of 20 µl at 450 – 500 ng µl^-1^ of each sample plasmid was then used as input into a Wheat germ TNT® high yield SP6 kit (Promega L3260) and incubated for 2 h at 25°C. A empty vector HALO tag control plasmid was also expressed to correct for non-specific HALO-tag linker DNA-binding in downstream analyses. All samples and control were performed in triplicate.

For each sample, 20 µl of Magne Halo Tag Beads (Promega G7281) were washed 3x and resuspended in 85% of the original volume in wash buffer (1x phosphate buffered saline and 0.05% NP-40). To bind TFs to magnetic beads 50 µl of wash buffer was added to each tube followed by 40 µl of TNT® Wheat Germ expression reaction for each sample, the remaining 10 µl of TNT® Wheat Germ expression reaction was retained for Western Blotting. Suspensions were then incubated for 1 h on a rotator at room temperature. Suspensions were then placed on a magnetic rack to precipitate the protein/magnetic bead complexes, then the supernatant was removed. Samples were then removed from the magnetic rack and the beads were resuspended in 100 µl of wash buffer and allowed to sediment at the bottom of the tube before placing on the magnetic rack again, beads were then washed 4 more times and then resuspended in 40 µl of wash buffer. Then 100 ng of adapter ligated DNA library was added to each tube in addition to 10 µg of salmon sperm DNA (Sigma Aldrich D9156) to reduce non-specific background protein DNA interactions. Volumes were then made up to 100 µl using Buffer EB (Qiagen 19086) and then incubated on a rotor for 1 h at room temperature. Samples were then placed on a magnetic rack and the DNA/protein/bead complexes were allowed to precipitate and the supernatant removed. The DNA/protein/beads were then removed from the rack and resuspended in 100 µl of wash buffer allowed to sediment at the bottom of the tube, then placed on the magnetic rack and allowed to precipitate before removal of the supernatant. DNA/protein/beads were then washed 4 more times and transferred to a new tubes for the final wash step. DNA/protein/beads were then resuspended in 16 µl of Buffer EB and incubated for 10 mins at 98°C then immediately placed on ice. Samples were then place a magnetic rack and beads were allowed to precipitate. 15 µl of the supernatant was then used as input for DNA-library indexing in the NEBNext Ultra II DNA library prep kit (New England Biolabs inc. E7645S) using a total of 14 cycles for PCR amplification, following the manufacturers protocol with minor modification: following the final clean up DAP-seq libraries were eluted in 12 µl of buffer EB and allowed to elute for 10 mins. DAP-seq libraries were then inspected on a 4150 Agilent Tapestation system using D1000 screen tapes (Agilent) and quantified with a Qubit™ 4 Fluorometer (Thermo Fisher Scientific) using Qubit™ dsDNA Quantitation Broad Range Assay (Thermo Fisher Scientific Q32850). Individual DAP-seq libraries were then normalised and pooled. Sequencing for DAP-seq libraries was carried out on a Illumina NovaSeq X 10B flow cell using 2x 150 bp paired-end sequencing, targeting 30,000,000 read pairs per sample.

### Western blot

Western blotting was carried out in parallel with DAP-seq library production. From the retained 10 µl of TNT High Yield Wheat Germ Expression reactions, 1 µl of each sample was mixed with 2x Lamelli Sample Buffer (Bio-Rad 1610737) and incubated at 90°C for 5 min. Then 0.5 µl was loaded onto 15% Criterion™ TGX Stain Free Protein Gels (Bio-Rad 5678085) and 4 µl of Precision Plus Protein Dual Color Standards (Bio-Rad 1610374) and run at 100 V for 90 min. Proteins were transferred to nitrocellulose membranes using a Trans-Blot Turbo Transfer System (Bio-Rad) according to the manufacturers instructions. Membranes were blocked in 5% (w/v) skim milk in TBS for 1 hr with gentle rocking then incubated at 4°C overnight with 1/2500 Anti-Halo Tag® Monoclonal Antibody (Promega G9211) in 0.1% TBST with gentle rocking. Membrane was then washed 3 x 10 min in 0.1% TBST then incubated with 1/20,000 Anti-Mouse IgG HRP Conjugate antibody (Promega W4021) in 0.1% TBST for 1 h at room temperature with gentle rocking. Membrane was then washed 3 x 20 min with 0.1% TBST. Membrane was then incubated for 2 min with SuperSignal™ West Pico PLUS Chemiluminescent Substrate (Thermo Fischer Scientific 34580) and imaged using a ChemiDoc MP Imaging System (Bio-Rad Laboratories, Hercules, CA, USA) using the colorimetric setting with 0.1 s exposure to capture the ladder, then Chemi setting with 0.1 s exposure to capture proteins, followed by merging of the images for final blot image.

### DAP-seq analysis

Fastq files were quality-filtered and adapters were trimmed using Trim Galore (v.0.6.10) with parameters -q 20, --illumina, and --trim-n. Paired-end reads were processed in paired end mode, and read quality was assessed using FastQC then aligned to the cs10 reference genome (GCF_900626175.2) using bowtie2 (v. 2.4.5) using default parameters^147^. Output BAM files were then filtered using samtools (v. 1.16.1) view with parameters -q 10 -F 2820, sorted using samtools sort, and index using samtools index^148^. Control BAM replicates were merged using samtools merge command. Peaks were called using MACS2 (v. 2.2.7.1) callpeak function with a relaxed cut-off threshold of --pvalue 1e-2 to introduce sufficient noise for downstream irreproducible discovery rate (IDR) analysis, with parameters --keep-dup all, --call-summits, -f BAMPE, and -g 736579359 using the merged control BAM file as INPUT control (Supplementary Table 5b-e)^123^. The fraction of reads in peaks (FRiP) was calculated for each replicate and the top two performing replicates were taken forward for IDR^149^. To determine high-quality reproducible peaks we used IDR (v. 2.0.4.2) with parameters --rank p.value and -i 0.05 on peak replicates sorted by -log10(p-value)^73^. To determine DAP-seq success we analysed the fraction of reads in IDR peaks where successful TFs contained ≥ 0.05 FRiP scores (Supplementary Table 5f).

As DAP-seq is performed using naked gDNA *in vitro* a large proportion of observed binding events are biological false positives in the context of chromatin therefore, to remove false positive binding events we filtered DAP-seq peaks using accesible chromatin peaks from the trichome lineage atlas. For a binding event to be considered real at least 50% of each DAP-seq peak must have been overlapped by an ATAC-seq peak from the trichome lineage peak set, this threshold fraction of 50% ensures that the centre of the DAP-seq peak, where the true TF binding site lays, is accessible *in vivo*. *De novo* motif discovery was performed using BaMM-motif with a second-order Bayesian Markov model (order = 2) and a motif length of 10 bp^150^. Motifs were identified from accessible DAP-seq peak sequences, using both DNA strands with a second-order background model. Motif occurrences were scanned using a p-value threshold of 1 x 10^-4^, and identified motifs were compared internally against the JASPAR2018 CORE database (Supplementary Table 5h-k)^151^. For validation, motif discovery was performed on those ATAC-set peaks that overlapped accessible DAP-seq peaks.

To identify trichome-lineage cell types enriched for accesible DAP-seq binding events, we quantified a per-cell chromatin enrichment score for each transcription factor using the corresponding DAP-seq peak set projected onto the single-nucleus ATAC assay. Enrichment scores were extracted for all cells in the trichome-lineage atlas and compared across annotated cell types. For each cell type, enrichment scores were tested against the pooled distribution of all other cells using a two-sided Wilcoxon rank-sum test in the Presto (v. 1.0.0) R-package, and resulting P values were adjusted for multiple testing using the Benjamini-Hochberg (BH) procedure. The magnitude of the enrichment was quantified using the rank-biserial estimate for the Wilcoxon test. Cell types with significantly (adjusted *P* < 0.05) were interpreted as enriched for accessible binding events

Genes associated with accessible DAP-seq binding events were identified by intersecting accessible DAP-seq peaks with regulatory regions derived from the cs10 (GCF_900626175.2) genome annotation, including the promoter region defined as 2 kb upstream of transcription start sites and first introns. Genes containing accessible DAP-seq peaks within these regions were retained and intersected with genes detected in the trichome-lineage scRNA-seq data. Per-cell enrichment scores for these genes were calculated using the Seurat AddModuleScore function, an implementation of the Tirosh module scoring method introduced in Seurat v2^152,153^. Enrichment scores were displayed as boxplots grouped by annotated cell type, with mean values overlaid as points. Statistical significance for each cell type was assessed against the pooled cellular background using a two-sided Wilcoxon rank-sum test followed by Benjamini-Hochberg correction, and the magnitude of enrichment was quantified using the rank-biserial correlation. Significance annotations were plotted only for cell types with adjusted *P* values <0.05 and median enrichment scores exceeding the overall median.

We implemented scMultiMap to identify enhancer-gene interactions in the trichome lineage (Supplementary Table 5u)^154^. To this end we tested for correlation between genes expressed in the trichome lineage (29,675) and accessible peaks in the trichome lineage (36485), with a maximum +/- 500 Kbp distance of interaction. Peak-gene enhancer correlations were then filtered for strong and significant correlations (cor > 0.5, adjusted *P* < 0.05). Neck and disc cell program DAP-seq associated peaks were then individually filtered against the scMultiMap results to determine genes linked with neck and disc cell program TFs.

## Supporting information

All_Supplemental_Tables

## Data availability

Raw sequencing reads for single nucleus ATAC and RNA libraries, as well as DAP-seq libraries were submitted to the NCBI sequencing read archive under BioProject ID PRJNA1457084. A ShinyCell2 web-based multiomic cell atlas browser and data analysis tool was made available for public use at https://singlecellcannabis.latrobe.edu.au/. Nuclei isolation and DAP-seq protocols were made available at protocols.io and are cited in the references.

## Code availability

All custom analysis scripts used in this study are available at GitHub: https://github.com/leecon30/cannabis-trichome-multiome-analysis. The repository contains scripts for single-nucleus multiome processing, quality control, cell-type annotation, trajectory inference, motif accessibility analysis, DAP-seq peak processing, peak–gene association analysis and figure generation.

## Acknowledgements

We thank Benji Wakely and Saksham Arora for their assistance with high-performance computing infrastructure, including support for deploying and serving the Shiny application associated with this study. MGL acknowledges Lucas Auroux for discussions on methodology. Work in the Lewsey lab is funded by the Australian Research Council Industrial Transformation Hub in Medicinal Agriculture (*IH180100006*) and Centre of Excellence in Plants for Space (*CE230100015*).

## Contributions

L.J.C., A.B., and M.G.L. conceived of and planned the project. L.J.C. maintained plants, isolated nuclei, prepared 10X snMultiome sequencing libraries, prepared DAP-seq sequencing libraries, and performed all formal data analysis. L.J.C. and L.C.L. contributed to RNA *in situ* hybridization. M.T.W. performed Micro-computed tomography and prepared images for publication. L.J.C and J.H. contributed to western blotting of DAP-seq proteins. L.J.C. and M.G.L. interpreted the results and drafted the manuscript. A.B. and M.T.W. provided critical feedback on manuscript.

## Ethics declarations

### Competing interests

The authors declare no competing interests.

## Extended data

**Extended Data Fig. 1.**
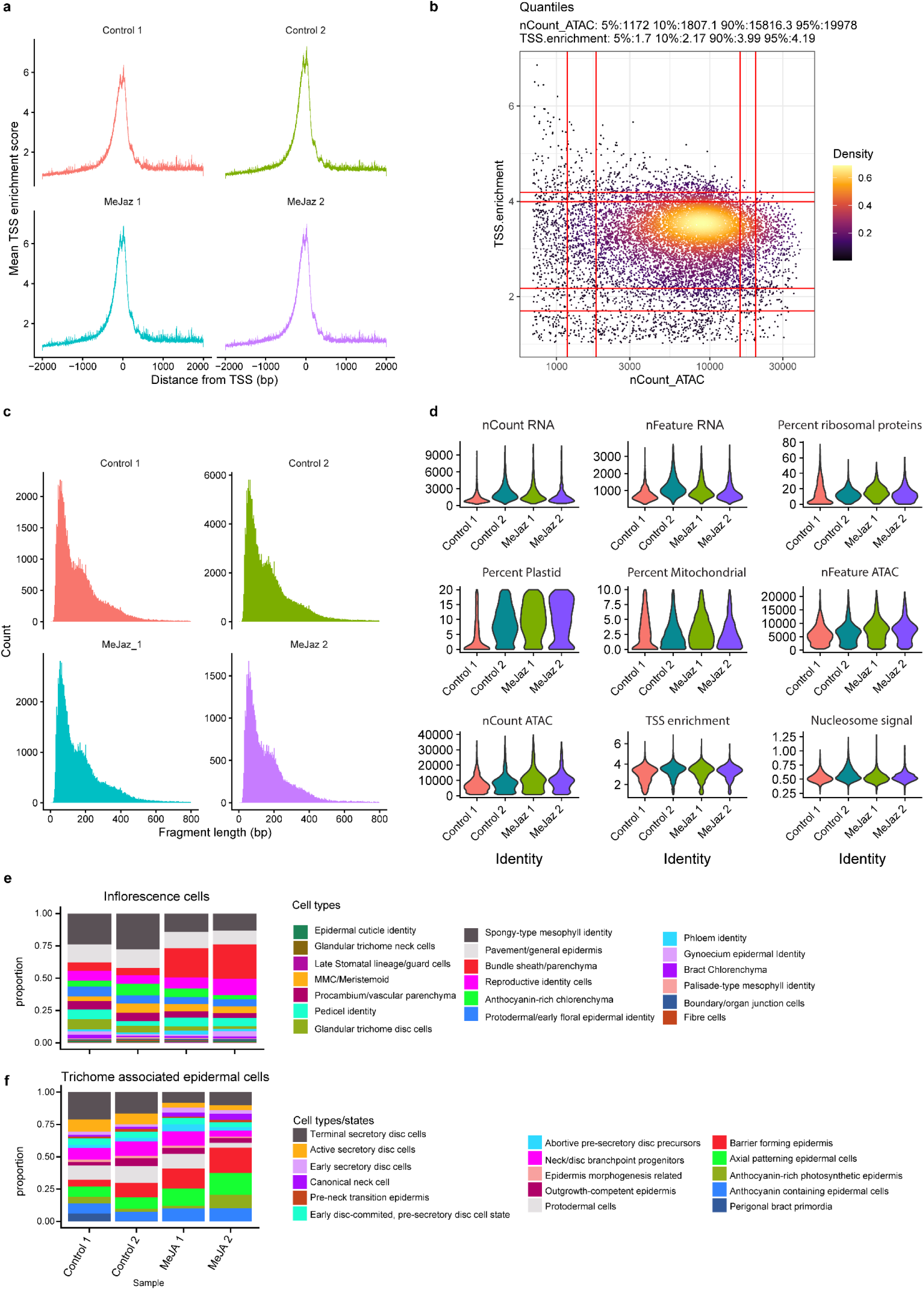
| Quality control of *Cannabis sativa* inflorescence single-nucleus multiome data. **a**, Transcription start site (TSS) enrichment profiles for the ATAC modality of each of the four samples after quality-control filtering. **b,** Density scatter plot of TSS enrichment score versus ATAC counts per nucleus for filtered inflorescence cells. **c,** Fragment size distribution of ATAC fragments from filtered inflorescence cells, showing a prominent nucleosome-free peak at approximately 90 bp and a periodic pattern consistent with mono- and di-nucleosomal fragments. **d**, Violin plots showing sample-level quality-control metrics. Metrics shown are RNA counts per nucleus (nCountRNA), detected genes per nucleus (nFeature RNA), percentage of ribosomal protein transcripts, percentage of plastid-derived transcripts (Percent Plastid), percentage of mitochondrial-derived transcripts (Percent Mitochondrial), detected ATAC peaks per nucleus (nFeature ATAC), ATAC counts per nucleus (nCount ATAC), TSS enrichment score and nucleosomal signal. **e**, Proportional stacked bar plots showing cell-type composition of the global inflorescence atlas for each sample. **f**, Proportional stacked bar plots showing cell-type composition of the trichome associated epidermal atlas for each sample.

**Extended Data Fig. 2.**
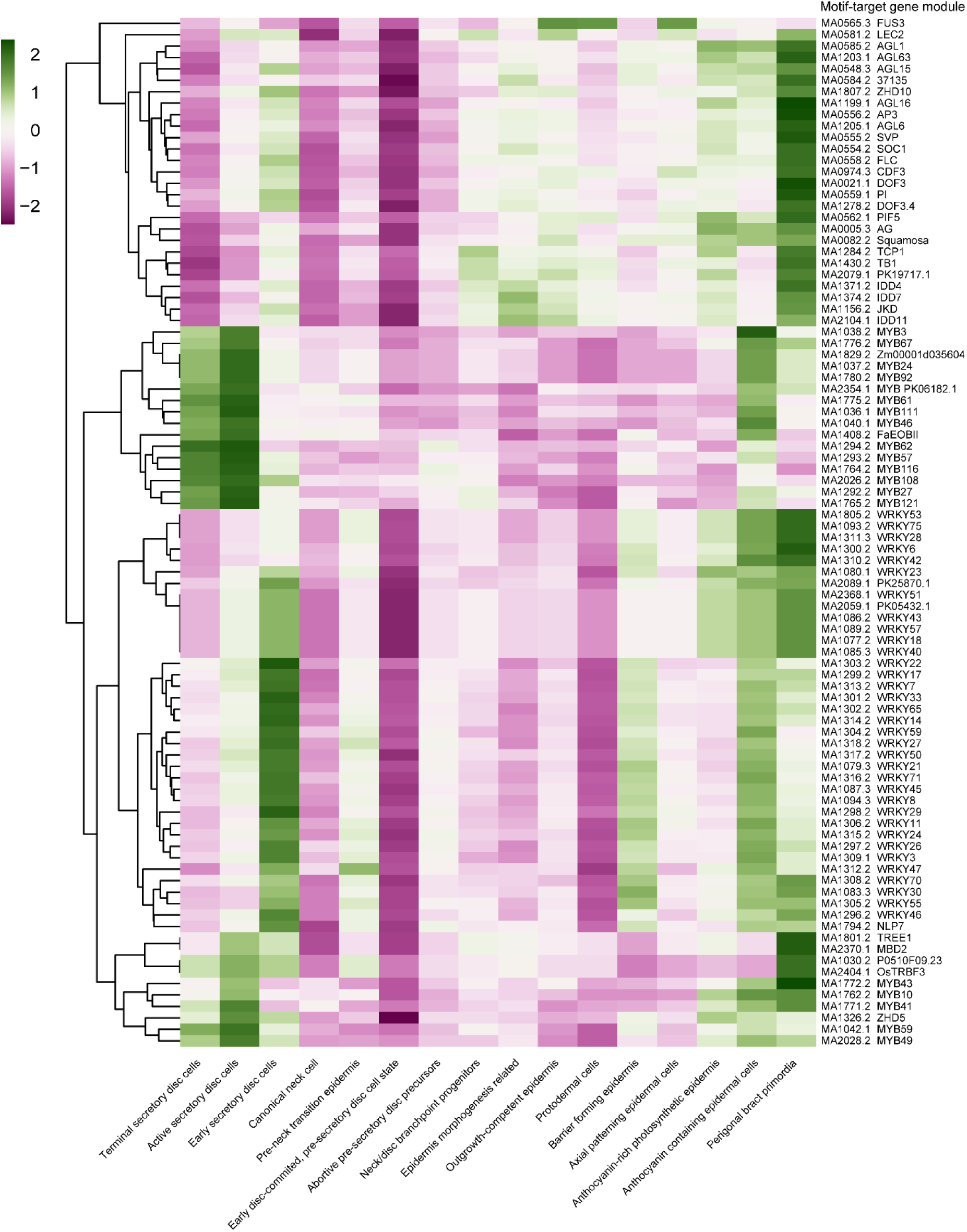
| Cell-type distribution of motif-associated target-gene module scores in the trichome-associated epidermal atlas. Heatmap showing motif target-gene module enrichment scores across cell types in the trichome-associated epidermal atlas. JASPAR 2024 motifs showing significant cell-type-enriched accessibility by chromVAR (AUC = 0.75, logFC > 1.5, adjusted *P* < 0.05) were used to define target-gene modules, which were subsequently scored across cell types. Green denotes higher module scores while red denotes lower module scores.

**Extended Data Fig. 3.**
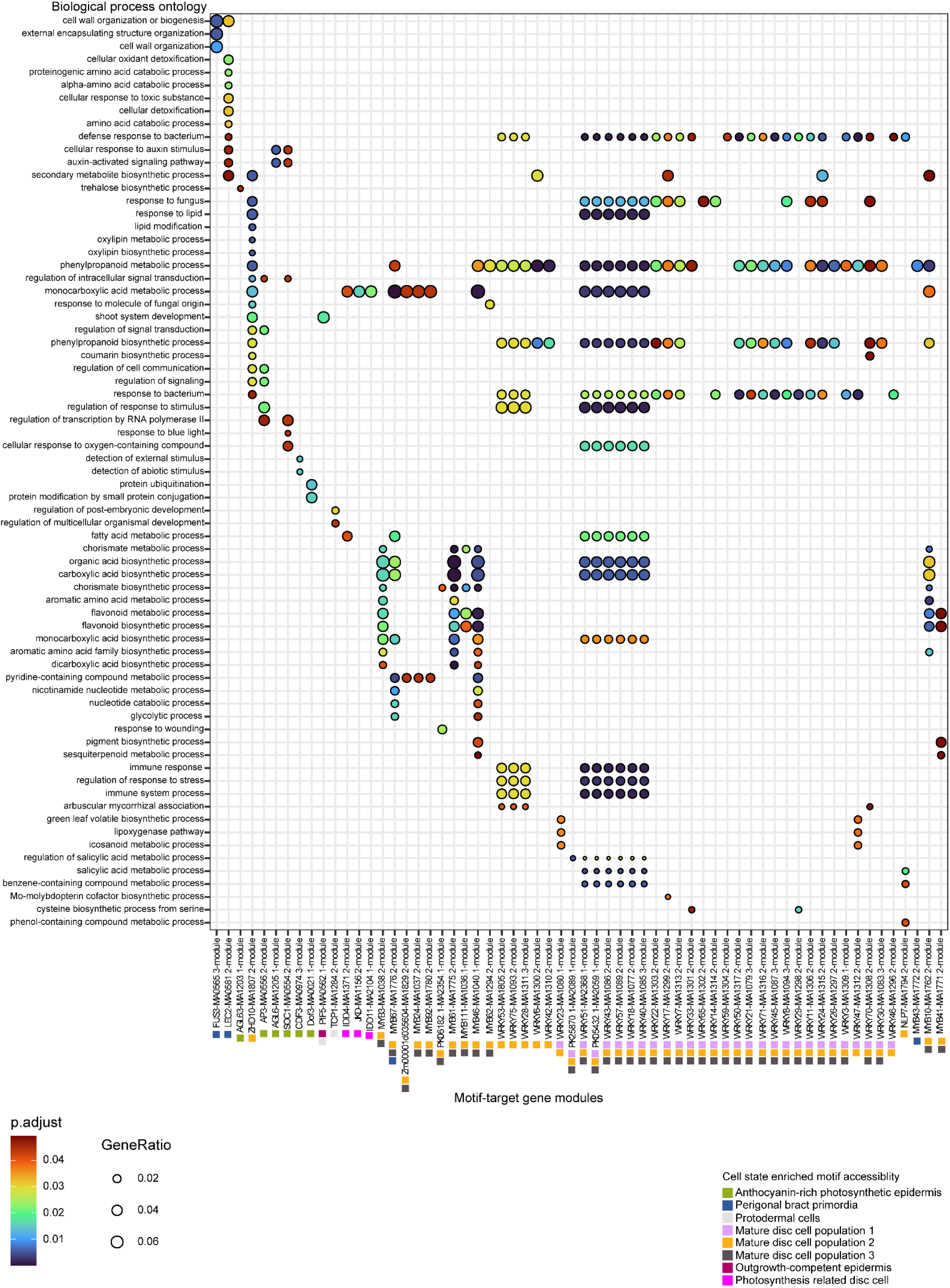
| Gene ontology biological process enrichment of motif-associated target-gene modules. Dot plot showing GO biological process enrichment for motif -associated target-gene modules in the trichome-associated epidermal atlas. Motifs from JASPAR 2024 showing significant cell-type-enriched accessibility by ChromVAR (AUC > 0.75, logFC > 1.5, adjusted *P* < 0.05) were used to define target-gene modules, which were then analysed for GO biological process enrichment using *clusterProfiler* with cs10 PANNZER2 annotations. Small colour-coded squares beneath each motif label on the x axis indicate the cell type in which motif accessibility was enriched, as defined in the legend. Dot size indicates gene ratio and dot colour indicates adjusted *P* value.

**Extended Data Fig. 4.**
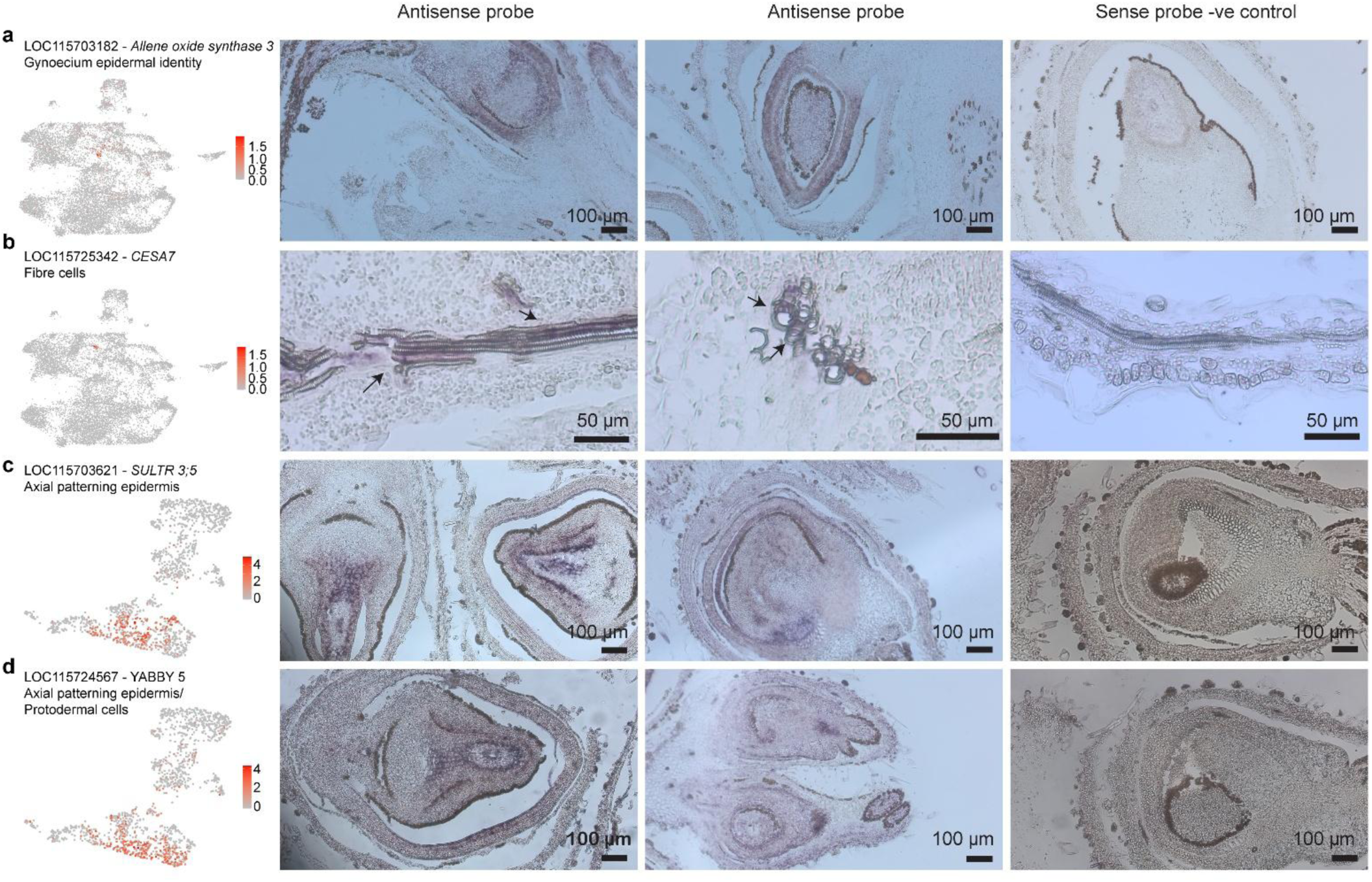
| RNA *in situ* hybridization. RNA *in situ* hybridization on sections of 5-week post floral induction (PFI) *C. sativa* inflorescences was used to validate cell-type assignments in the inflorescence and trichome-associated epidermal atlas. Antisense probe signal (purple) was interpreted relative to the corresponding sense-probe negative controls. **a**, *AOS3* (LOC115703182) transcripts in gynoecium epidermal identity cells. Signal specifically marks gynoecium epidermal cells along the carpel and stigma. **b**, *CESA7* (LOC115725342) transcripts in the vascular bundle. Signal is restricted to vascular strand containing helically thickened xylem elements. **c**, *SULTR 3;5* (LOC115703621) transcripts in axial patterning epidermis. Signal was restricted to developing flower epidermis layers. **d**, *YABBY5* (LOC115724567) transcripts mark axial patterning epidermis and protodermal cells. Signal marks early axial patterning epidermis and is also observed in protodermal cells in developing flowers.

**Extended Data Fig. 5.**
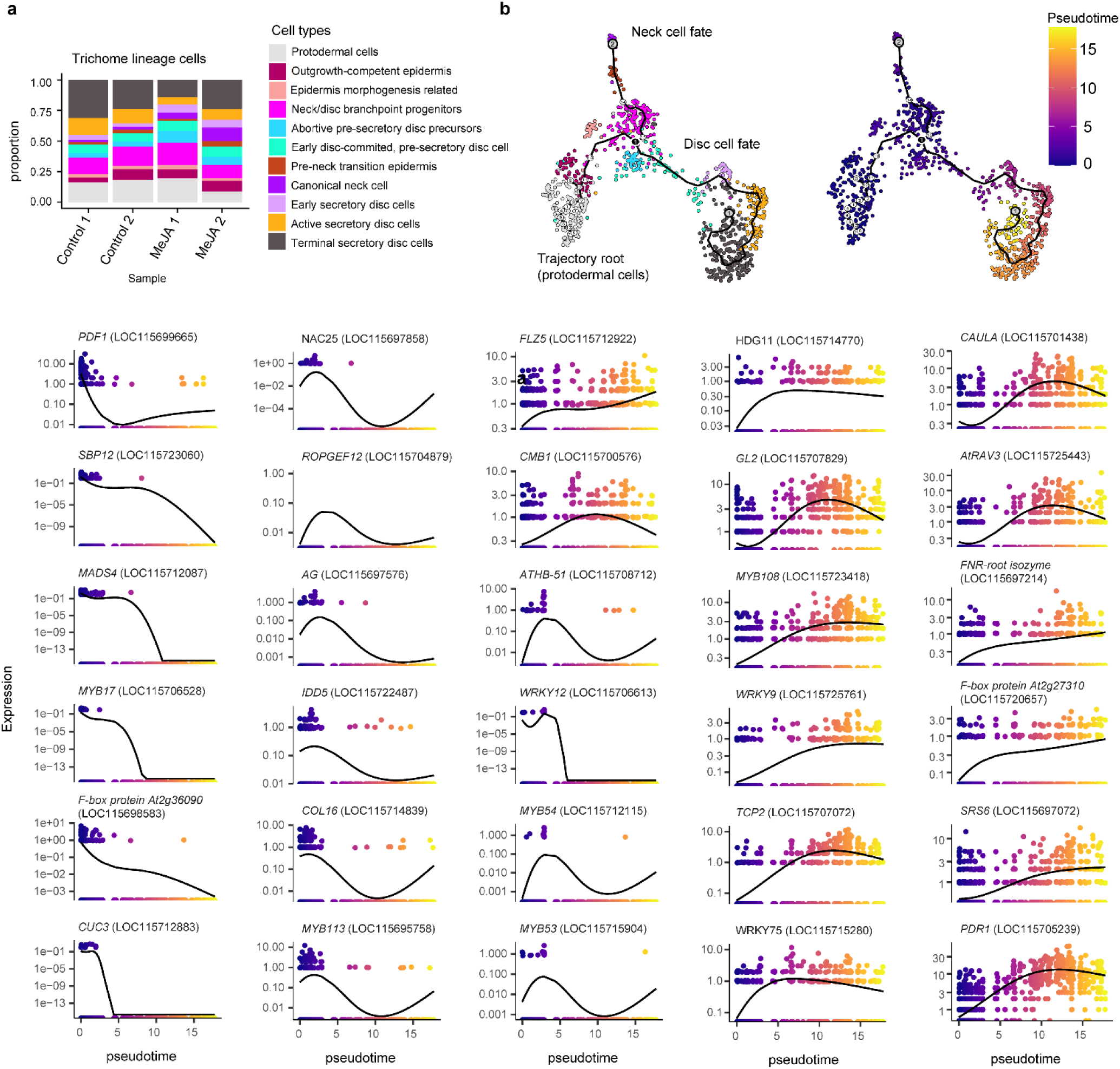
| Transcription factors with pseudotime-structured expression in the trichome lineage. **a**, Proportional stacked bar plots showing cell-type composition of the trichome-associated epidermal cell atlas for each sample. **b**, Smoothed expression profiles of transcription factors with trajectory-structured expression in the trichome lineage, plotted across pseudotime. Transcription factors were identified on the basis of significant spatial autocorrelation along the trajectory (Moran’s *I* > 0.01, adjusted *P <* 0.05) and selected from early and late pseudotime intervals.

**Extended Data Fig. 6.**
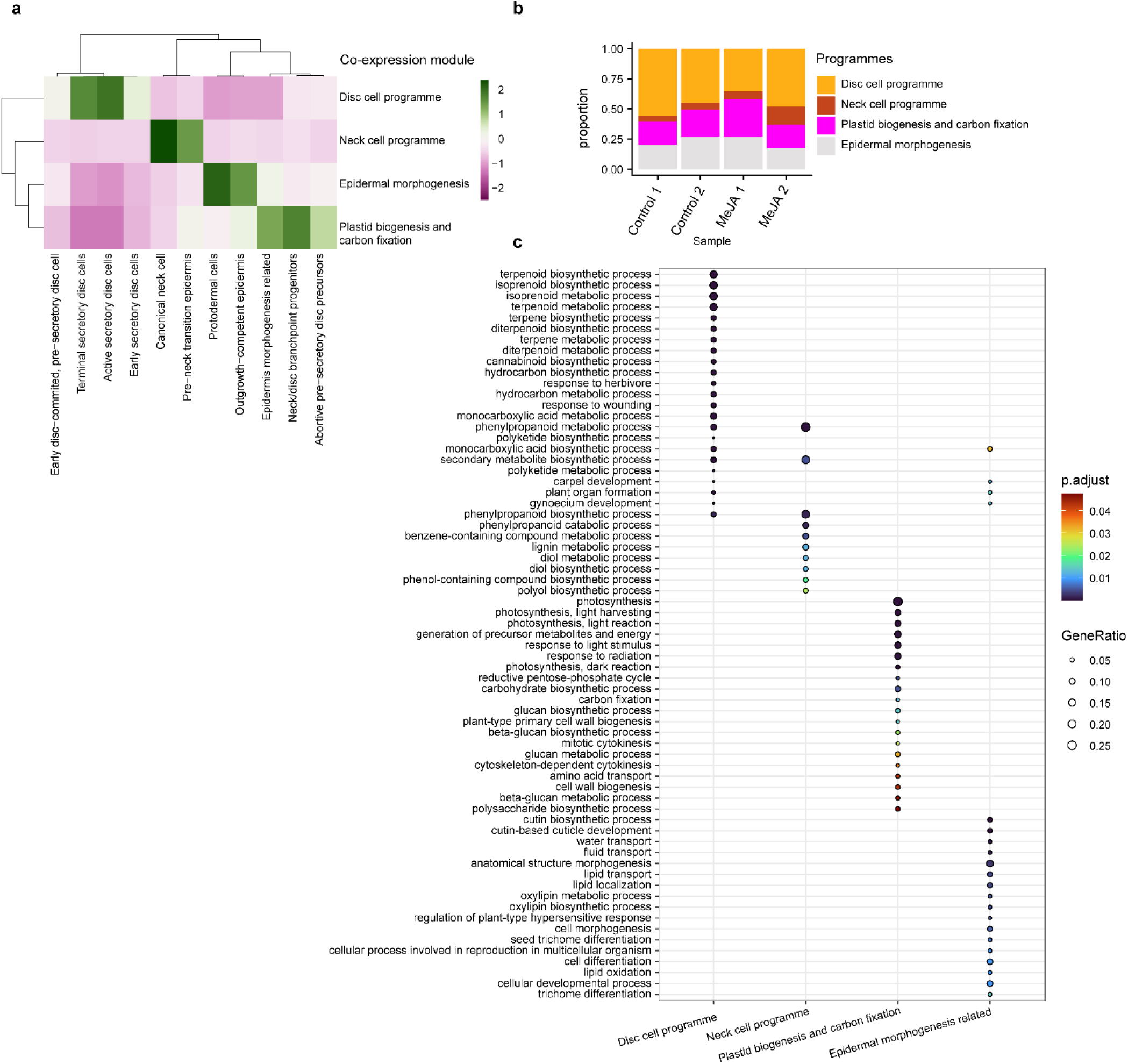
| Trajectory guided gene co-expression modules reveal regulatory programmes. **a,** Heatmap showing enrichment scores of trajectory-guided gene co-expression modules across cells in the trichome lineage. Gene modules were identified using Monocle3 from genes exhibiting significant trajectory-structured expression (Moran’s *I* > 0.10, *q* < 0.01). Module-enriched cells were defined on the basis of aggregated module expression (threshold > 1), and genes were retained within each module when mean expression across module-enriched cells exceeded 0.5). **b,** Proportional stacked bar plots showing cell-type composition of the trichome lineage atlas for each sample. **c**, Dot plot showing GO biological process enrichment of trajectory-guided co-expression modules. Disc cell programme module associated with disc cell states was enriched for specialised metabolism terms, neck cell programme module was enriched for lignin- and phenylpropanoid-related processes, plastid biogenesis and carbon fixation module was enriched for photosynthesis-related terms, and epidermal morphogenesis modules were enriched for trichome differentiation and developmental processes.

**Extended Data Fig. 7.**
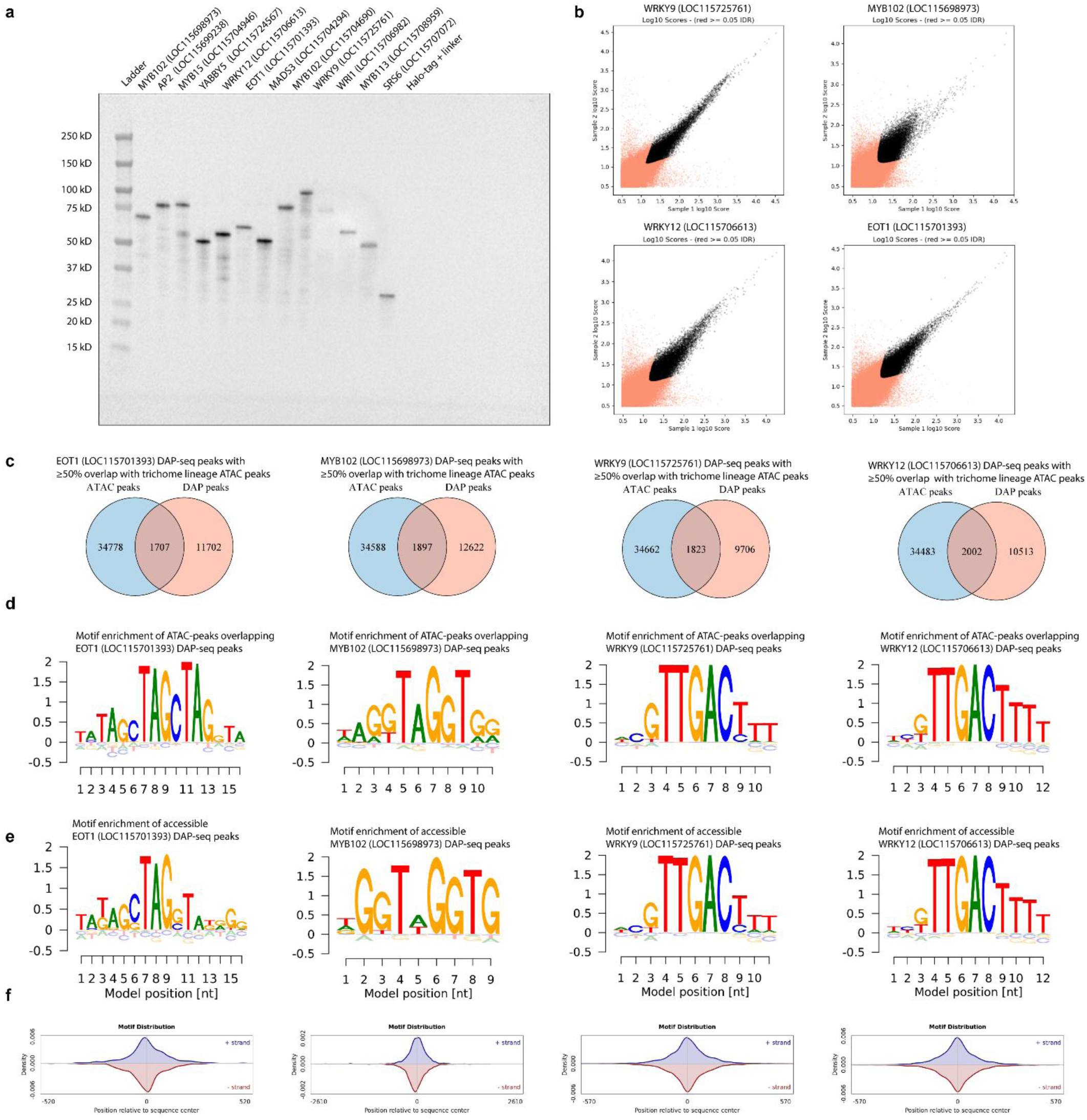
| Quality control of DNA affinity purification assay. **a**, Western blot of trichome-lineage-associated transcription factor proteins generated using the TnT® SP6 High Yield Wheat Germ Protein Expression System. Proteins were detected with Promega Anti-Halo primary antibody and an HRP-conjugated anti-mouse secondary antibody. Molecular masses were estimated relative to Precision Plus Protein Dual Colour Standards. **b**, IDR concordance plots for DAP-seq replicates of WRKY9, MYB102, WRKY12 and EOT1, showing log10 peak scores in replicate 1 versus replicate 2. Black points indicate peaks passing the IDR threshold, whereas red points indicate peaks failing the threshold. All four transcription factors showed reproducible peak sets with concordant signal between replicates. **c,** Venn diagrams showing overlap between trichome lineage ATAC peaks and DAP-seq peaks for EOT1, MYB102, WRKY9 and WRKY12. Peaks were considered overlapping when a trichome lineage ATAC peak covered at least 50% of the DAP-seq peak. Numbers indicate unique and shared peaks. **d,** Sequence logos showing motifs enriched by BAMMmotif in trichome lineage ATAC peak intervals overlapping DAP-seq peaks for EOT1, MYB102, WRKY9 and WRKY12. **e**, Sequence logos of motifs enriched by BAMMmotif in DAP-seq peaks overlapping trichome lineage ATAC peaks for EOT1, MYB102, WRKY9 and WRKY12. **f**, Motif distribution plots for EOT1, MYB102, WRKY9 and WRKY12, showing the positional density of motif occurrences relative to the centre of accessible DAP-seq peak intervals. Blue and red curves indicate motif density on the positive and negative strands, respectively. Motif occurrences were centrally enriched in all four datasets.

**Extended Data Fig. 8.**
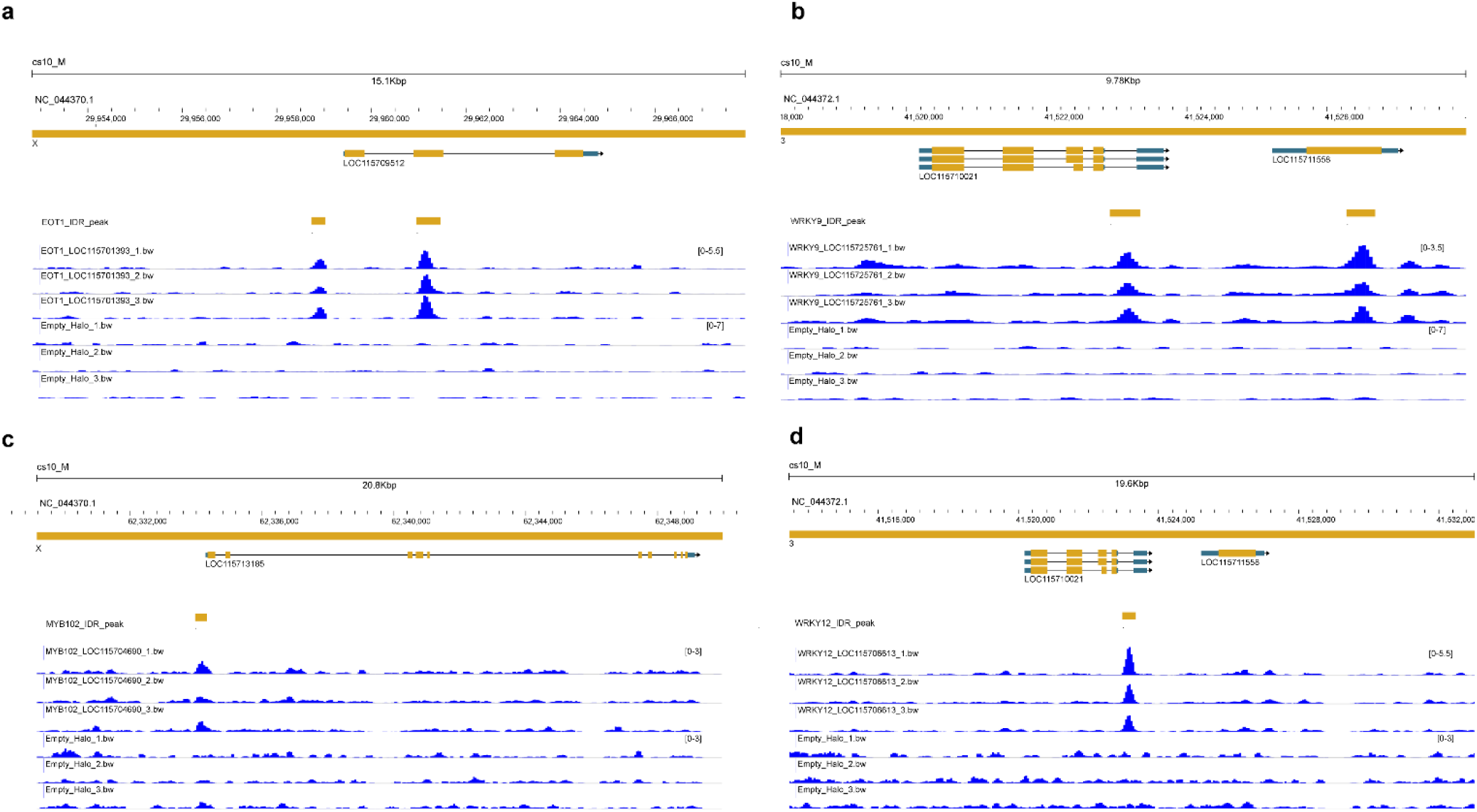
| Genome browser views support reproducible DAP-seq peak enrichment over empty HALO controls. **a–d**, JBrowse2 views of representative DAP-seq peak regions for **a**, EOT1, **b**, WRKY9, **c**, MYB102 and **d**, WRKY12. For each transcription factor, three biological replicate coverage tracks are shown together with three empty HALO control tracks. Gene models and IDR peak calls are displayed above the signal tracks, illustrating reproducible enrichment above control background at representative bound loci.

**Extended Data Fig. 9.**
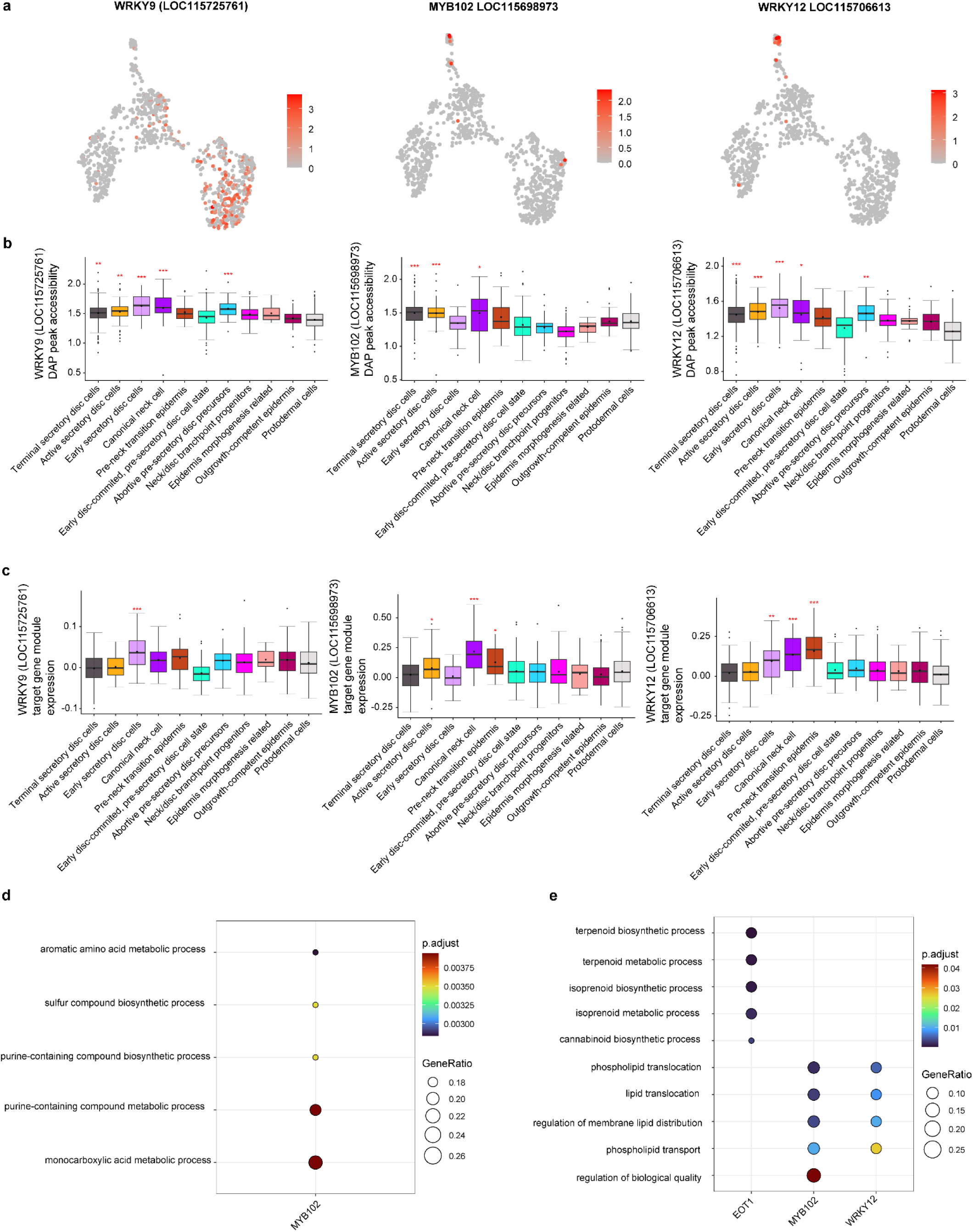
| Functional enrichment of DAP-seq-supported target gene sets. **a**, UMAP of trichome-lineage cells showing *WRKY9* (LOC115725761), *MYB102* (LOC115698973), *WRKY12* (LOC115706613) expression. **b**, Boxplots showing enrichment of accessible chromatin peaks containing *WRKY9*, *MYB102*, and *WRKY12* binding motifs across-trichome lineage cell types. significance was assessed using two-sided Wilcoxon tests with adjusted *P* values (* < 0.05, ** < 0.01, *** < 0.0001). **c**, Boxplots showing expression scores of WRKY9, MYB102, and WRKY12 motif-associated target-gene modules across trichome-lineage cell types. Target genes were inferred from accessible peaks containing motifs in disc cell programme cells for WRKY9 and neck cell programme cells for MYB102 and WRKY12. significance was assessed using two-sided Wilcoxon tests with adjusted *P* values (* < 0.05, ** < 0.01, *** < 0.0001). **d**, Dot plot showing GO biological process enrichment of motif-associated target-gene modules, with significant enrichment detected only for the MYB102 module. GO analysis was performed using cs10 PANZZER2 annotations. **e**, Dot plot showing GO biological process enrichment of genes linked by scMultiMap to accessible DAP-seq peaks for EOT1, MYB102 and WRKY12. Linked genes were identified using significant peak–gene associations passing the specified correlation (cor > 0.5) and adjusted *P* value thresholds, and GO analysis was performed using cs10 PANZZER2 annotations.

## Supplementary information

## Notes

### Competing Interest Statement

The authors have declared no competing interest.

